# The Impact of Species Tree Estimation Error on Cophylogenetic Reconstruction

**DOI:** 10.1101/2023.01.24.525446

**Authors:** Julia Zheng, Yuya Nishida, Alicja Okrasińska, Gregory M. Bonito, Elizabeth A.C. Heath-Heckman, Kevin J. Liu

## Abstract

Just as a phylogeny encodes the evolutionary relationships among a group of organisms, a cophylogeny represents the coevolutionary relationships among symbiotic partners. Both are widely used to investigate a range of topics in evolutionary biology and beyond. Both are also primarily reconstructed using computational analysis of biomolecular sequence data as well as other biological character data. The most widely used cophylogenetic reconstruction methods utilize an important simplifying assumption: species phylogenies for each set of coevolved taxa are required as input and assumed to be correct. Many theoretical and experimental studies have shown that this assumption is rarely – if ever – satisfied, and the consequences for cophylogenetic studies are poorly understood. To address this gap, we conduct a comprehensive performance study that quantifies the relationship between species tree estimation error and downstream cophylogenetic estimation accuracy. The study includes performance benchmarking using *in silico* model-based simulations. Our investigation also includes assessments of cophylogenetic reproducibility using genomic sequence datasets sampled from two important models of symbiosis: soil-associated fungi and their endosymbiotic bacteria, and bobtail squid and their bioluminescent bacterial symbionts. Our findings conclusively demonstrate the major impact that upstream phylogenetic estimation error has on downstream cophylogenetic reconstruction quality.

## Introduction

A cophylogeny represents the evolutionary and coevolutionary relationships among multiple sets of coevolved taxa, and cophylogenies are widely used to study fundamental and applied topics throughout biology and the life sciences [Blasco-Costa et al., 2021, Martínez-Aquino, 2016]. For example, untangling coevolutionary histories is essential to reconstructing the web of life [Thompson, 2010], as symbiosis and coevolution has played an important role in evolution at different scales – from genes to proteins, biomolecular pathways, organisms, populations, and beyond [Libeskind-Hadas et al., 2014].

As is the case in phylogenetic estimation, cophylogenies are principally reconstructed using computational analyses of biomolecular sequences as well as other types of biological data [Dismukes et al., 2022]. The most widely used computational approach for cophylogenetic estimation consists of a multi-stage pipeline where: (1) a species tree is independently estimated for each coevolved set of taxa using the same approaches as in a traditional phylogenetic study, and (2) a cophylogeny is then estimated using the estimated species trees as input, alongside the known host and symbiont associations. Next-generation biomolecular sequencing technologies have transformed phylogenetics and our broader understanding of evolutionary biology [Czech et al., 2022], and there exists great interest in the scientific community to use cophylogenetic methods to help understand ancient and recent coevolution of symbiotic species (Figure 1).

**Fig. 1.**
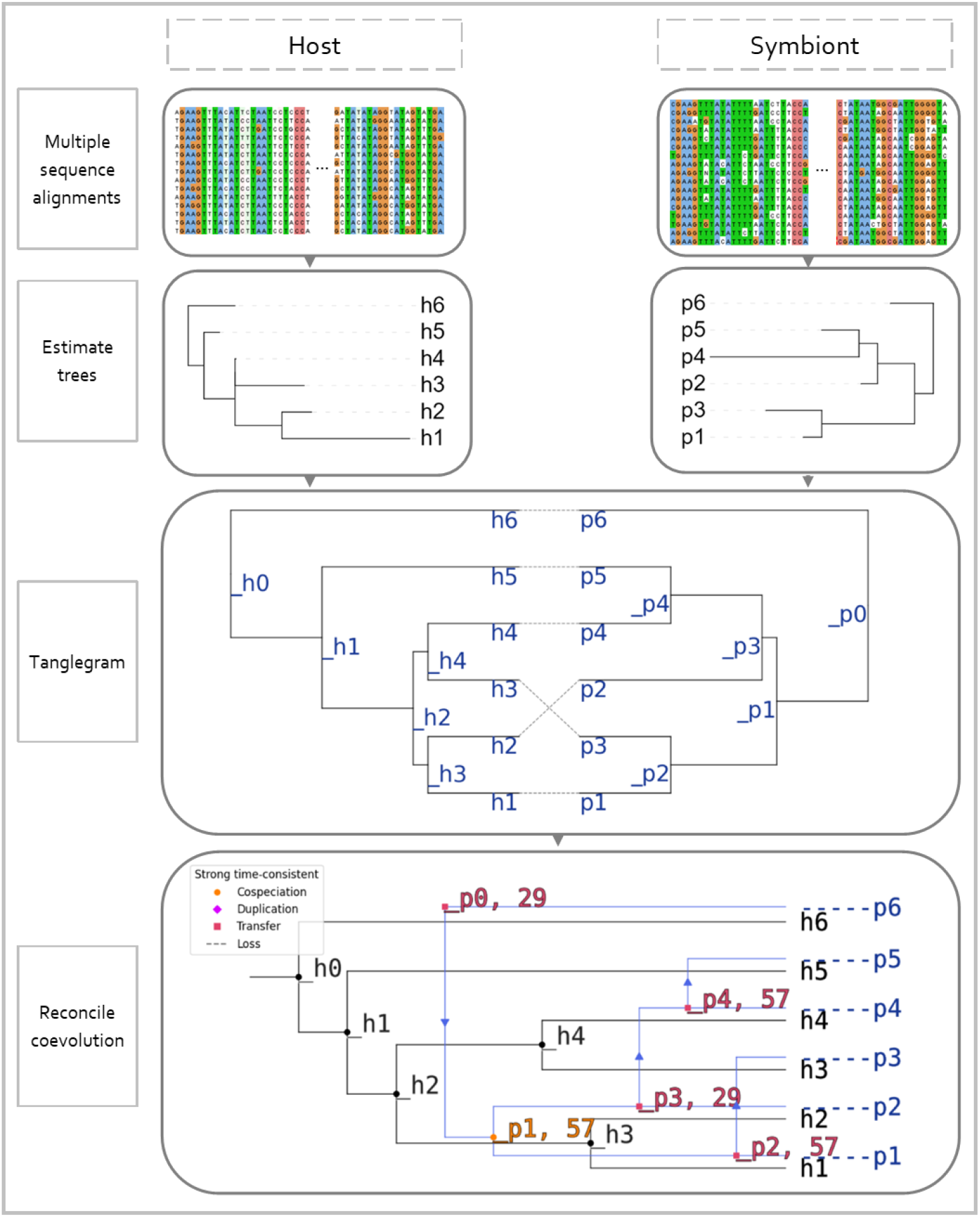
A typical workflow for cophylogenetic reconstruction. (1) Biomolecular sequence data for host taxa and symbiont taxa are aligned. (2) A species tree is independently estimated using each multiple sequence alignment as input. (3) The tanglegram corresponding to the estimated host tree, estimated symbiont tree, and known host/symbiont associations is produced. (4) Finally, a cophylogeny is reconstructed using the tanglegram as input. The cophylogeny maps topological structure in the host tree to corresponding topological structure in the symbiont tree based on shared coevolutionary history, where each relation in the mapping corresponds to a coevolutionary event (e.g., a cospeciation event, a host-switching event, etc.). Example dataset from [Hafner et al., 1994].

Many cophylogenetic methods have been developed and they fall into two broad categories: (1) statistical tests of overall congruence between host and symbiont tree topologies, such as PARAFIT [Legendre et al., 2002], PACo [Balbuena et al., 2013], and MRCAlink [Schardl et al., 2008], and (2) event-based methods that perform phylogenetic reconciliation using either parsimony-based optimization or, less commonly, model-based statistical optimization. EMPRess [Santichaivekin et al., 2021], Jane [Conow et al., 2010], Treemap [Charleston and Page, 2002], COALA [Baudet et al., 2015], and CoRe-PA [Merkle et al., 2010] are examples of event-based methods. Event-based methods typically account for multiple types of coevolutionary events [Charleston, 1998]: cospeciation (or codivergence or codifferentiation) involving both host and symbiont lineages, duplication of a symbiont lineage within a host lineage, loss of a symbiont lineage within a host lineage, and host shift (or host switch) where a symbiont lineage’s association switches to a different host lineage. In this study, we focus on event-based cophylogenetic reconstruction methods to investigate a finer granularity of evolutionary and coevolutionary event reconstructions.

The multi-stage pipeline design requires a critically important assumption: the estimated species trees in the first stage are used directly in the second stage under the assumption that they are correct. However, it is well understood in traditional phylogenetics that many factors can cause phylogenetic estimation methods to return some degree of estimation error, and estimation errors introduced in upstream computational tasks are important factors to consider. For example, numerous studies have investigated the strong impact that upstream multiple sequence alignment error can have on subsequent gene tree estimation [Liu et al., 2010]. But this insight conflicts with the prevailing assumption made by cophylogenetic reconstruction pipelines. Contributing to this oversight is the lack of similar studies investigating this issue directly [Dismukes et al., 2022].

To address this gap, we have undertaken a study to examine the relationship between upstream phylogenetic estimation error and downstream cophylogeny reconstruction accuracy. Our performance study utilizes both simulated and empirical datasets that span a range of evolutionary conditions, and we validate and quantify the major impact that the former has on the latter.

## Methods

Our performance study included a comprehensive suite of simulated benchmarking datasets that spanned a range of evolutionary conditions. The simulation conditions differed in terms of number of taxa, sequence length, evolutionary divergence, and distribution of coevolutionary event types. Figure 2 provides an illustrated overview of the simulation study procedures.

**Fig. 2.**
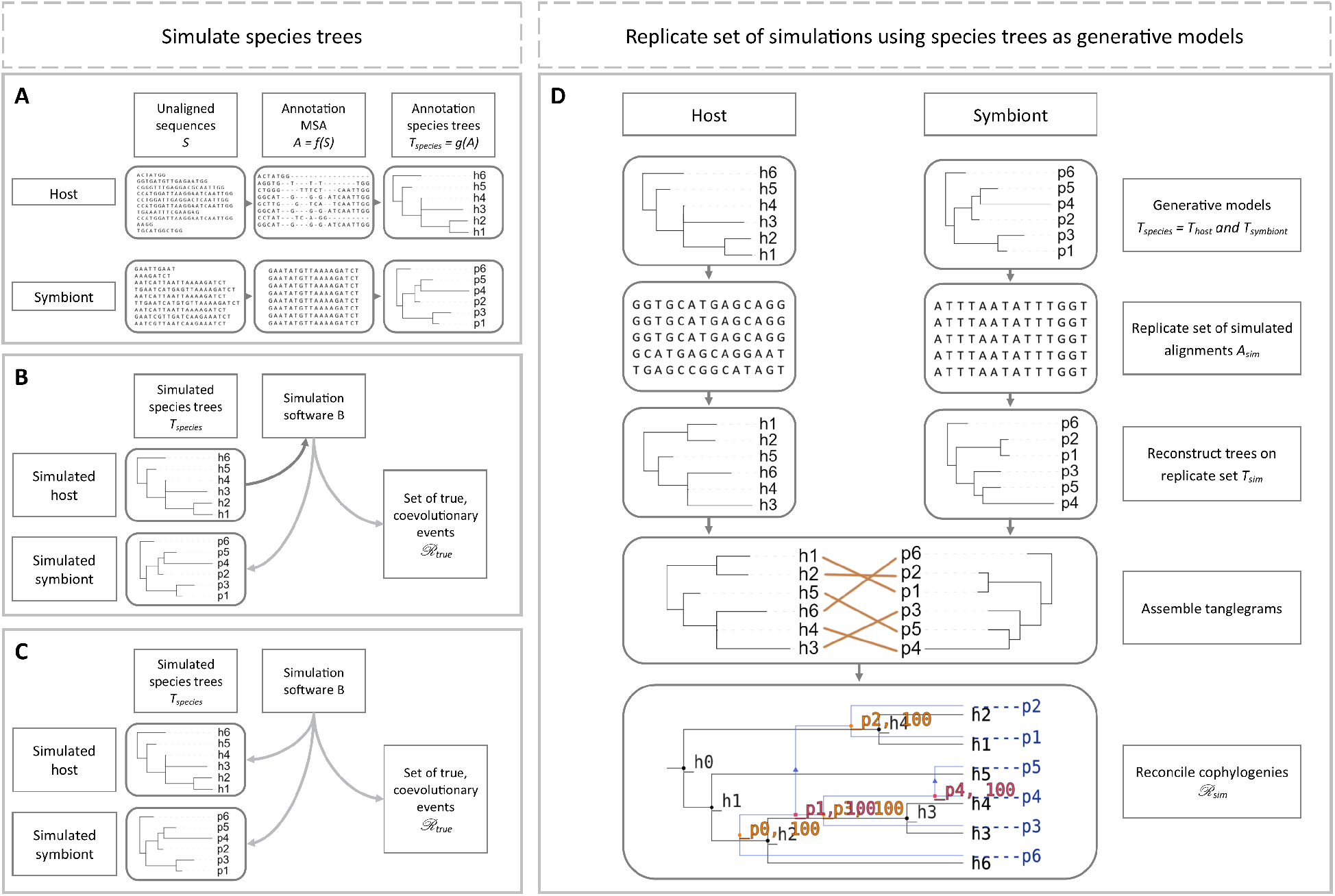
Illustrated overview of simulation study experiments. Three simulation procedures were used to simulate datasets. The procedures differed in the cophylogeny model and simulation software that they utilized. (A) The “mixed” simulations utilized model cophylogenies and constituent species trees that were based on empirical dataset analyses. (B) The “backward-time” simulations sampled model cophylogenies under the backward-time model of [Avino et al., 2019]. (C) The “forward-time” simulations sampled model cophylogenies under Treeducken’s forward-time model [Dismukes and Heath, 2021]. (D) For each model cophylogeny, sequence evolution along each constituent species tree was simulated under finite-sites models, resulting in a multiple sequence alignment. The simulation procedure was repeated to obtain *k* experimental replicates. Once the simulation procedure has concluded, phylogenetic and cophylogenetic reconstruction is performed using a computational pipeline. For each replicate dataset, a phylogenetic tree is reconstructed for host taxa using their corresponding multiple sequence alignment as input, and similarly for symbionts. The estimated host tree and estimated symbiont tree are combined with host/symbiont association data to produce a tanglegram. The tanglegram is then used as input to reconstruct a cophylogeny.

The simulation experiments utilized one of three different simulation procedures. First, the “mixed” simulations utilized an empirically estimated cophylogeny and its constituent species trees and host/symbiont associations as the phylogenetic models for *in silico* simulation of biomolecular sequence evolution. Second, the “backward-time” simulations were conducted using the backward-time cophylogeny model of [Avino et al., 2019]. Third, a fully *in silico* set of simulations were run using the forward-time cophylogeny model proposed by [Dismukes and Heath, 2021], which we refer to as the “forward-time” simulations. Cophylogenetic and phylogenetic method performance on each simulated dataset was then assessed with respect to reference or ground truth.

We also performed comparative analyses of two empirical genomic sequence datasets. One empirical dataset consists of cephalopod hosts and their bacterial symbionts, which serve as a well-studied model of open symbiosis (i.e., partnerships arising from horizontal transmission between hosts and/or the environment); the other dataset was sampled from fungal hosts and their bacterial endosymbionts, which are an emerging model of closed symbiosis (i.e., partnerships whose coevolution involves strictly vertical descent over time). The two systems thus provide a comparative contrast along a spectrum of symbiotic partnership flexibility [Perreau and Moran, 2022].

### Definitions

We now introduce mathematical background needed to describe the experimental procedures. Some of the notation and definitions follow [Wieseke et al., 2015].

A rooted phylogenetic tree 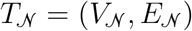 is a rooted evolutionary history for a set of taxa 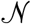. We note that many cophylogenetic reconstruction algorithms require rooted binary phylogenetic trees as input. The rooted binary tree 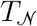 has a root *ρ* with in-degree zero and out-degree two, leaves 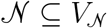 where each leaf has out-degree zero and in-degree one, and inner nodes 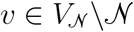 where each inner node has out-degree two and in-degree one. For each directed edge 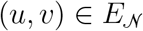, *v* is a child of *u*. Each edge is also denoted by *e_v_* with branch length *u* 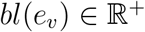. For vertices *u, v* ∈ *V_n_, u* is an ancestor of *v, u* ∈ *anc*(*v*), *v* is a descendent of *u*, and *u* ∈ *desc*(*v*) if and only if *u* lies on the unique path from root *ρ* to *v*.

For a pair of rooted phylogenetic trees *T_H_* and *T_S_* denoting the evolutionary history of a set *H* of hosts and a set *S* of symbionts, respectively, *T_H_* is the host tree and *T_S_* is the symbiont tree. A mapping function *ϕ*(*s, h*): *S* × *H* → {0, 1} denotes known interactions between the extant species of *T_H_* and *T_S_*, where *ϕ*(*s, h*) = 1 means a symbiont is associated with a host, and otherwise *ϕ*(*s, h*) = 0. The set (*T_H_, T_S_, ϕ*) is called a tanglegram and serves as the input to cophylogenetic methods. A cophylogenetic reconciliation or reconstruction is defined as the set of event associations 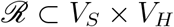 between the internal nodes of the symbiont tree *T_S_* and the internal nodes of the host tree *T_S_*. For a symbiont *s*, an event association 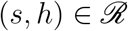 means *h* is one of the host species known to have been associated with *s*.

The unrooted version 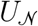 of a rooted phylogenetic tree 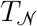 can be obtained by converting all directed edges into undirected edges, deleting the root, and connecting its incident edges into a single remaining edge. Equivalently, an unrooted binary tree 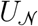 on the leaf set 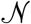 has internal nodes with degree three and leaves with degree one, and each leaf represents a distinct taxon in the taxon set 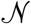.

Tree topology differences were evaluated with normalized Robinson-Fould (nRF) distances. Given an unrooted tree U, a bipartition is created by removing an edge from *U* to generate two subtrees *t*_1_ and *t*_2_, where trivial bipartitions are defined as a subtree containing only a leaf node. For two unrooted trees *U*_1_ and *U*_2_ with the same set of leaf nodes 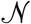, the non-trivial bipartitions are given by *B*_1_ and *B*_2_, respectively. The Robinson-Fould (RF) metric is the cardinality of the symmetric difference between the sets of non-trivial bipartitions that appear in *T*_1_ and *T*_2_, which is |*B*_1_ – *B*_2_| + |*B*_2_ – *B*_1_|. The normalized RF distance is calculated by dividing RF distance by the maximum RF distance between two trees with *n* taxa, which is 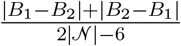.

Reconciled cophylogenetic events were statistically evaluated with a calculation from existing literature [Wieseke et al., 2015] defined as follows. Let 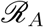 and 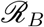 be the reconstructed event associations of all internal vertices from cophylogenetic reconciliation of tanglegram A and tanglegram B, respectively. Then, the proportion of reconciled events in 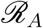 that were also found in 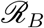 is 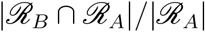.

### Simulation study

#### Mixed simulations

Six empirical datasets were obtained from literature, from single-locus datasets with sequence length under 1 kb to next-generation-sequencing (NGS) multi-locus datasets with sequence length well over 1 Mb (Table 1). The sequence data were preprocessed and aligned using MAFFT v7.221 with default settings [Katoh and Standley, 2013]. Species phylogenies were reconstructed from concatenated multiple sequence alignments under the General Time Reversible (GTR) model of nucleotide substitution with Γ model of rate heterogeneity [Yang, 1996] and midpoint rooted using RAxML v8.2.12 [Stamatakis, 2014]. Some of the cophylogenetic reconstruction methods under study were limited to one-to-one host/symbiont associations; symbiont taxa were subsampled as needed to address this limitation. Cophylogenetic events were estimated with eMPRess [Santichaivekin et al., 2021] from the host and symbiont phylogenies and host-symbiont associations.

**Table 1.** Summary statistics for mixed simulation datasets. Each mixed simulation condition (“Model conditions”) is based on a previously published cophylogenetic study (“Source”). For each dataset type (either host or symbiont, as denoted by “Taxa”), the number of taxa (“# taxa”), true MSA length (“Aln length”), average and standard error of normalized Hamming distance of true MSAs (“ANHD Avg” and “ANHD SE”, respectively), and average and standard error of model tree height (“Height Avg” and “Height SE”, respectively) are reported. The number of cospeciation, duplication, host switch, and loss events in the reference cophylogeny are reported as “# cospec, “# dup”, “# switch”, and “# loss”, respectively.

| Model conditions | Source | Taxa | # taxa | Aln length | ANHD Avg | ANHD SE | Height Avg | Height SE | # cospec | # dup | # switch | # loss |
| --- | --- | --- | --- | --- | --- | --- | --- | --- | --- | --- | --- | --- |
| mixed-gopher | [Hafner et al., 1994] | Host | 15 | 379 | 0.2241 | 0.0007 | 0.4024 | 0.0042 | 9-10 | NA | NA | NA |
|  |  | Symbiont | 17 | 379 | 0.5249 | 0.0007 | 3.0598 | 0.0359 |  |  |  |  |
| mixed-stinkbug | [Hosokawa et al., 2006] | Host | 7 | 1,745 | 0.2371 | 0.0016 | 0.2651 | 0.0016 | 6 | NA | NA | NA |
|  |  | Symbiont | 12 | 1,583 | 0.0661 | 0.0006 | 0.1349 | 0.0011 |  |  |  |  |
| mixed-primate | [Switzer et al., 2005] | Host | 55 | 696 | 0.2599 | 0.0002 | 0.6079 | 0.0046 | 22 | NA | NA | NA |
|  |  | Symbiont | 41 | 425 | 0.3376 | 0.0004 | 0.8169 | 0.0050 |  |  |  |  |
| mixed-damselfly | [Lorenzo-Carballa et al., 2019] | Host | 24 | 1,051 | 0.1734 | 0.0004 | 0.4919 | 0.0036 | 5 | 7 | 10 | 40 |
|  |  | Symbiont | 23 | 3,297 | 0.1327 | 0.0004 | 0.2643 | 0.0010 |  |  |  |  |
| mixed-moth | [Zhang et al., 2014] | Host | 82 | 1,404 | 0.1021 | 0.0001 | 0.2147 | 0.0013 | 14-28 | 20-28 | 5-10 | 74-106 |
|  |  | Symbiont | 53 | 4,326 | 0.0250 | 0.0000 | 0.0486 | 0.0003 |  |  |  |  |
| mixed-bird | [de Moya et al., 2019] | Host | 37 | 5,000 | 0.1087 | 0.0001 | 0.1526 | 0.0009 | 12 | NA | 4 | NA |
|  |  | Symbiont | 57 | 5,000 | 0.3562 | 0.0001 | 0.5459 | 0.0011 |  |  |  |  |

The empirical estimate for each dataset (specifically the constituent species phylogenies and continuous parameter values which are associated with the model cophylogeny) served as the statistical model for downstream *in silico* simulation. The reconstructed species trees (including branch lengths and other continuous parameter estimates) served as generative models from which multiple sequence alignments were simulated using Seq-Gen [Rambaut and Grass, 1997].

We also performed two additional simulation experiments to investigate the impact of evolutionary divergence and sequence length. In simulations with varying evolutionary divergence, model tree branch lengths were multiplied by a scaling parameter *h*. We explored a range of settings for the parameter *h* where each set of experiments selected a setting from the set {0.1, 0.5, 1, 2, 5, 10}. The simulations with varying sequence length were based on the mixed-bird model condition, where simulated sequence length was reduced from over 1 Mb to 5 kb.

#### Backward-time simulations

The backward-time model of [Avino et al., 2019] was used to simulate coevolution among *n* host taxa and *n* symbiont taxa, as well as host/symbiont associations. Our simulations explored varying numbers of taxa *n* ∈ {10, 50, 100, 500}. The simulations made use of a custom-modified Python program that was originally implemented by Avino et al. [2019] (Table 2). The simulation program takes a host tree as input and simulates a symbiont tree backward-in-time along the host tree by randomly drawing wait times to determine the timing and type of coevolutionary event(s) on a particular host tree branch. We used INDELible to sample host trees under a random birth-death model (see Supplementary Materials for more details). Model trees were deviated away from ultrametricity using Moret et al. [2002]’s approach with deviation factor *c* = 2.0 [Nelesen et al., 2007]. We used custom scripts to perform the ultrametricity deviation calculations. We note that the Avino et al. [2019]’s simulation software does not directly provide the model cophylogeny as output. Instead, a reference cophylogeny was obtained using eMPRess estimation on the true model trees for host and symbiont taxa as input. The choice of reference cophylogeny allows comparison of cophylogenetic estimation when ground truth inputs are provided (i.e., true model trees) versus cophylogenetic estimation when estimated trees are used as input.

**Table 2.**
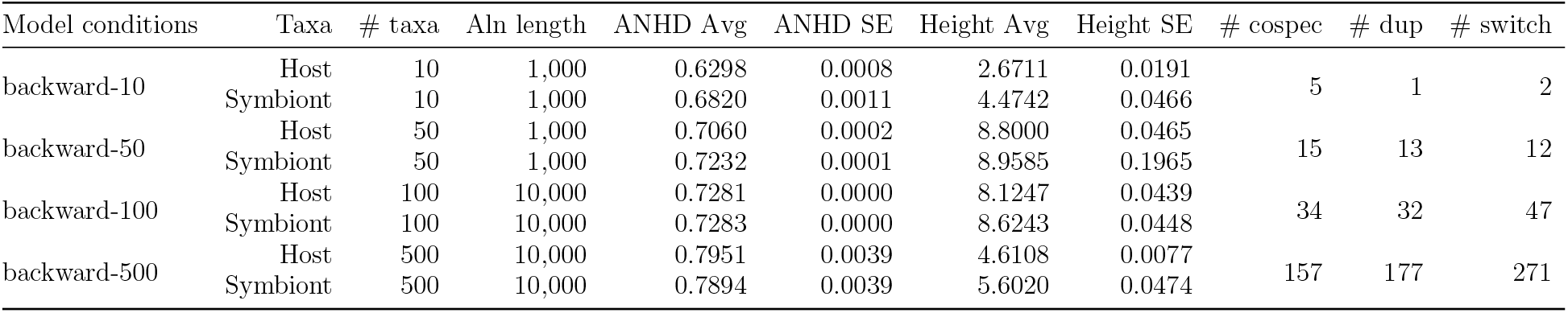
Summary statistics for backward-time simulation datasets. Each backward-time simulation condition (“Model conditions”) varied the number of host and symbiont taxa (“# taxa”) simulated under Avino et al. [2019]’s backward-time coevolutionary model. The simulations included cospeciation, duplication, and host switch events, but not loss events. Otherwise, table layout and description are identical to Table 1.

Simulation of sequence evolution along model phylogenies followed the same procedure as in the mixed simulations. The substitution model parameters were based on empirical estimates from our re-analysis of the dataset from [de Moya et al., 2019]’s study.

As with the mixed simulations, additional experiments with varying evolutionary divergence were performed using the backward-time simulation procedure. The scaling parameter *h* was similarly set to a value from {0.1, 0.5, 1, 2, 5, 10}.

#### Forward-time cophylogeny simulations

The forward-time simulations utilized Treeducken [Dismukes and Heath, 2021] and its forward-time coalescent model to sample a model cophylogeny (along with its associated species trees and host/symbiont associations). Model parameter settings (Table 3) were based on estimates from selected empirical datasets. The resulting five model conditions included a range of dataset sizes (i.e., number of taxa and sequence length), substitution rates, base frequency distributions, and coevolutionary event distributions (Table 4). Model tree branch lengths were deviated from ultrametricity using the same procedure as in the other simulation experiments.

**Table 3.** Treeducken parameters used in simulating forward-time datasets. Treeducken was used to simulate cophylogenies and their constituent species phylogenies under a forward-time coalescent-based model [Dismukes and Heath, 2021]. Treeducken’s model specifies the following parameters: the symbiont speciation rate *λ_S_*, the symbiont extinction rate *μ_S_*, the cospeciation rate *λ_C_*, the host speciation rate *λ_H_*, the host extinction rate *μ_H_*, the expected number of host taxa *H_tips_*, and the expected number of symbiont taxa *S_tips_*.

| Model condition | $H_{tips}$ | $S_{tips}$ | $\lambda_H$ | $\lambda_C$ | $\lambda_S$ | $\mu_H$ | $\mu_S$ | time |
| --- | --- | --- | --- | --- | --- | --- | --- | --- |
| forward-gopher | 35 | 55 | 0.3104 | 1.2000 | 0.0290 | 0 | 0 | 2.2 |
| forward-stinkbug | 35 | 55 | 0.2104 | 1.2000 | 0.0290 | 0 | 0 | 2.0 |
| forward-primate | 203 | 50 | 0.3374 | 0.6246 | 0.0452 | 0 | 0 | 4.8 |
| forward-damselfly | 25 | 25 | 0.1843 | 0.8846 | 0.2920 | 0 | 0 | 2.0 |
| forward-bird | 27 | 134 | 0.0544 | 0.6000 | 0.4520 | 0 | 0 | 4.0 |

**Table 4.** Summary statistics for forward-time simulation datasets. For each model condition (“Model conditions”), Treeducken was used to perform forward-time simulations based on a previously published cophylogenetic study (“Source”). Each simulated dataset consisted of a model cophylogeny, its constituent model species trees and host/symbiont associations, and true MSAs. Table layout and description are otherwise identical to Table 1.

| Model conditions | Source | Taxa | # taxa | Aln length | ANHD Avg | ANHD SE | Height Avg | Height SE | # cospec | # dup | # switch | # loss |
| --- | --- | --- | --- | --- | --- | --- | --- | --- | --- | --- | --- | --- |
| forward-gopher | [Hafner et al., 1994] | Host | 17 | 300 | 0.5664 | 0.0010 | 2.3260 | 0.0313 | 16 | 0 | 1 | 0 |
|  |  | Symbiont | 16 | 300 | 0.5426 | 0.0009 | 2.5639 | 0.0403 |  |  |  |  |
| forward-stinkbug | [Hosokawa et al., 2006] | Host | 16 | 1,000 | 0.5672 | 0.0012 | 4.2617 | 0.0707 | 14 | 0 | 2 | 0 |
|  |  | Symbiont | 14 | 1,000 | 0.5825 | 0.0016 | 3.9159 | 0.0326 |  |  |  |  |
| forward-primate | [Switzer et al., 2005] | Host | 48 | 400 | 0.6030 | 0.0002 | 8.0586 | 0.0791 | 31 | 3 | 17 | 0 |
|  |  | Symbiont | 34 | 400 | 0.7017 | 0.0004 | 10.7577 | 0.2931 |  |  |  |  |
| forward-damselfly | [Lorenzo-Carballa et al., 2019] | Host | 24 | 1,000 | 0.3437 | 0.0003 | 0.5804 | 0.0031 | 12 | 9 | 12 | 0 |
|  |  | Symbiont | 21 | 1,000 | 0.4233 | 0.0007 | 1.1334 | 0.0066 |  |  |  |  |
| forward-bird | [de Moya et al., 2019] | Host | 31 | 5,000 | 0.6953 | 0.0004 | 4.1329 | 0.0023 | 21 | 33 | 10 | 0 |
|  |  | Symbiont | 54 | 5,000 | 0.7125 | 0.0002 | 5.0964 | 0.0027 |  |  |  |  |

Additional experiments varying evolutionary divergence were performed with the forward-time simulation procedure, where the scaling parameter *h* was assigned a value from {0.1, 0.5, 1, 2, 5, 10}.

#### Experimental replication

For each model condition, the simulation procedure was repeated to obtain 100 replicate datasets. Results are reported across all replicate datasets in each model condition.

#### Phylogenetic and cophylogenetic reconstruction and assessment

On each simulated dataset, RAxML v8.2.12 was used to reconstruct a phylogenetic tree under the GTR model. Reconstructed phylogenies were midpoint rooted. The resulting phylogenetic estimates and host/symbiont associations were used by eMPRess [Santichaivekin et al., 2021] to perform cophylogenetic reconciliation using either default settings or alternative cophylogenetic event costs that were estimated using COALA [Baudet et al., 2015] and CoRe-PA [Merkle et al., 2010].

In each simulation study experiment, the topological error of an estimated tree was compared to its corresponding model tree based on normalized Robinson-Foulds distance. Each estimated cophylogeny was compared to either the model cophylogeny (in the case of the forward-time simulation experiments) or reference cophylogeny (in the case of the mixed and backward-time simulation experiments) based on [Wieseke et al., 2015]’s precision calculation.

### Empirical study of soil-associated fungi and their bacterial endosymbionts

#### Sample acquisition and sequencing

Isolates were collected and also sourced from established culture collections. Modified versions of the soil plate [Warcup, 1950] and selective-baiting method [Shirouzu et al., 2012] were used to isolate Mortierellomycotina from soil. The techniques described in [Bonito et al., 2016] were used to isolate Mortierellomycotina from pine and spruce roots.

In total, thirteen metagenomic samples of *Mortierella spp*. and their associated endobacteria were collected and sequenced (Table 5). Ten samples were sequenced using Illumina HiSeq 2500 short-read sequencing and three samples were sequenced using PacBio long-read sequencing.

**Table 5.**
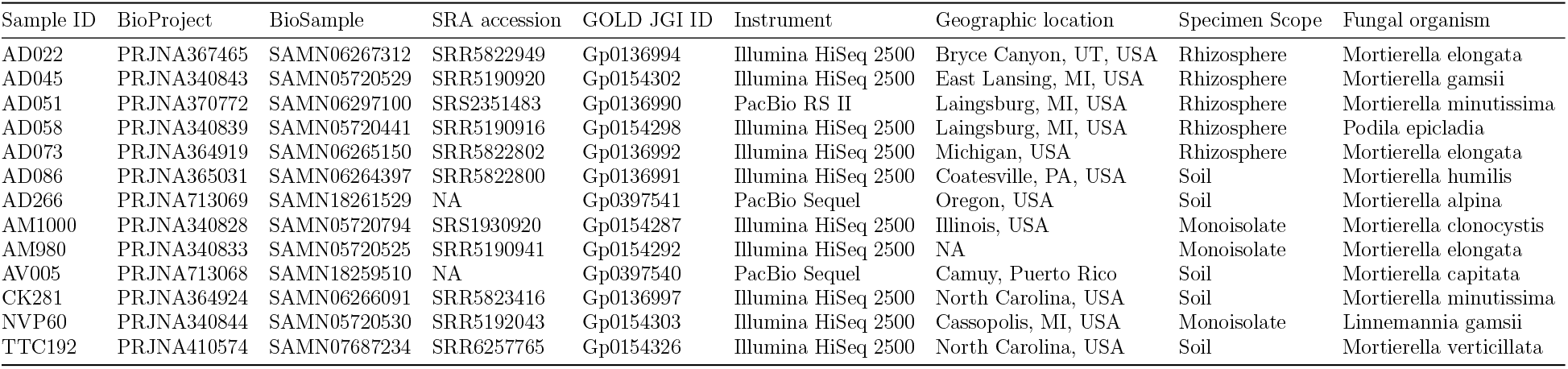
List of *Mortierella spp*. and endobacteria used in this study.

Illumina-sequenced metagenomic reads were trimmed with BBDuk (ftl=5 minlen=90) [Bushnell, 2018] to remove Illumina adapters, trim five leftmost bases, and discard reads shorter than 90 bp after trimming. The quality of trimmed reads was assessed by FastQC [Andrews, 2010]. De novo assembly of the metagenomic samples was conducted with SPAdes (-k 21,33,55,77,99,127) [Bankevich et al., 2012] to produce contigs. BBMap [Bushnell, 2018] was used to calculate summary statistics on assembled contigs. BUSCO [Simão et al., 2015] was used with the mucoromycota_odb10 and burkholderiales_odb10 databases to assess the completeness of de novo assembly and confirm the presence of endobacteria, respectively (Table 6).

**Table 6.**
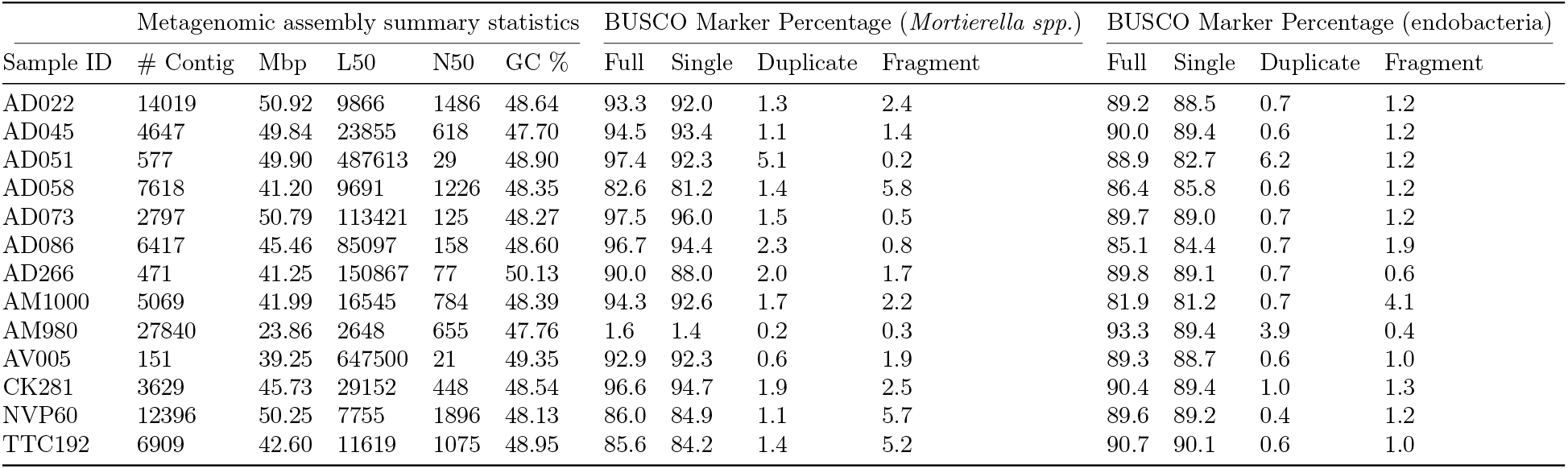
Summary statistics for *Mortierella spp*. and endobacterial assemblies.

| Sample ID | Metagenomic assembly summary statistics |  |  |  |  | BUSCO Marker Percentage (*Mortierella spp.*) |  |  |  | BUSCO Marker Percentage (endobacteria) |  |  |  |
| --- | --- | --- | --- | --- | --- | --- | --- | --- | --- | --- | --- | --- | --- |
|  | # Contig | Mbp | L50 | N50 | GC % | Full | Single | Duplicate | Fragment | Full | Single | Duplicate | Fragment |
| AD022 | 14019 | 50.92 | 9866 | 1486 | 48.64 | 93.3 | 92.0 | 1.3 | 2.4 | 89.2 | 88.5 | 0.7 | 1.2 |
| AD045 | 4647 | 49.84 | 23855 | 618 | 47.70 | 94.5 | 93.4 | 1.1 | 1.4 | 90.0 | 89.4 | 0.6 | 1.2 |
| AD051 | 577 | 49.90 | 487613 | 29 | 48.90 | 97.4 | 92.3 | 5.1 | 0.2 | 88.9 | 82.7 | 6.2 | 1.2 |
| AD058 | 7618 | 41.20 | 9691 | 1226 | 48.35 | 82.6 | 81.2 | 1.4 | 5.8 | 86.4 | 85.8 | 0.6 | 1.2 |
| AD073 | 2797 | 50.79 | 113421 | 125 | 48.27 | 97.5 | 96.0 | 1.5 | 0.5 | 89.7 | 89.0 | 0.7 | 1.2 |
| AD086 | 6417 | 45.46 | 85097 | 158 | 48.60 | 96.7 | 94.4 | 2.3 | 0.8 | 85.1 | 84.4 | 0.7 | 1.9 |
| AD266 | 471 | 41.25 | 150867 | 77 | 50.13 | 90.0 | 88.0 | 2.0 | 1.7 | 89.8 | 89.1 | 0.7 | 0.6 |
| AM1000 | 5069 | 41.99 | 16545 | 784 | 48.39 | 94.3 | 92.6 | 1.7 | 2.2 | 81.9 | 81.2 | 0.7 | 4.1 |
| AM980 | 27840 | 23.86 | 2648 | 655 | 47.76 | 1.6 | 1.4 | 0.2 | 0.3 | 93.3 | 89.4 | 3.9 | 0.4 |
| AV005 | 151 | 39.25 | 647500 | 21 | 49.35 | 92.9 | 92.3 | 0.6 | 1.9 | 89.3 | 88.7 | 0.6 | 1.0 |
| CK281 | 3629 | 45.73 | 29152 | 448 | 48.54 | 96.6 | 94.7 | 1.9 | 2.5 | 90.4 | 89.4 | 1.0 | 1.3 |
| NVP60 | 12396 | 50.25 | 7755 | 1896 | 48.13 | 86.0 | 84.9 | 1.1 | 5.7 | 89.6 | 89.2 | 0.4 | 1.2 |
| TTC192 | 6909 | 42.60 | 11619 | 1075 | 48.95 | 85.6 | 84.2 | 1.4 | 5.2 | 90.7 | 90.1 | 0.6 | 1.0 |

The PacBio-sequenced metagenomic reads were de novo assembled with CANU [Koren et al., 2017], with the exception of sample AV005: its draft assembly was obtained directly from JGI (Project ID: 1203140). Completeness and summary statistics were assessed in the same manner as for Illumina-sequenced assemblies (Table 6).

#### Variant calling

Fungal and endobacterial contigs were extracted from metagenomic assemblies and variants were called using one of three procedures, depending on the set of loci to be analyzed. Sequences with greater than 99.95% sequence similarity were pruned. The three resulting datasets consisted of: (1) all genomic loci, (2) CDS loci, and (3) rDNA genes. Summary statistics for each dataset are listed in Table 7.

**Table 7.** Summary statistics for processed *Mortierella spp*. and endobacterial MSAs. Alignment is abbreviated “Aln”, and average normalized Hamming distance is abbreviated “ANHD”.

| Model condition | Taxa | Summary statistics |  |  |  |
| --- | --- | --- | --- | --- | --- |
|  |  | # taxa | Aln length | Aln gappiness | Aln ANHD |
| full assembly | Fungus | 7 | 4,607,802 | 0.8194 | 0.0003 |
|  | Endobacteria | 7 | 215,165 | 0.4738 | 0.0022 |
| CDS genes | Fungus | 8 | 2,423,869 | 0.8337 | 0.0003 |
|  | Endobacteria | 8 | 152,860 | 0.5714 | 0.0013 |
| rDNA | Fungus | 5 | 87 | 0.6345 | 0.0041 |
|  | Endobacteria | 5 | 179 | 0.5218 | 0.0057 |

The all-genomic-loci dataset was processed using the following steps. Contigs were extracted using the draft genome *Linnemannia elongata* AD073 v1.0 (JGI Project ID: 1203123) as the reference genome for fungus and draft genome *Mycoavidus cysteinexigens* B1-EB (Genome ID: 1553431.3) from the PATRIC database as a reference for endobacteria; the reference fungal genome was processed using RepeatMasker [Chen, 2004]. BLASTN (-outfmt 6 -max_target_seqs 200) [Camacho et al., 2009] was used to identify fungus and endobacteria in the de novo assembly against the procured draft reference genome databases. Seqtk (subseq -l 60) [Li, 2018] analyzed BLAST hits to recover a draft fungal genome and a draft endobacteria genome from the de novo assembly. Variant calling was performed with the MUMmer package [Delcher et al., 2003] using the draft genomes against the reference genomes. Within the MUMmer suite [Delcher et al., 2003], NUCmer was used to align the draft genome against the reference and show-snps identified the single nucleotide variants (SNV). Then, the MUMmerSNPs2VCF software was used to convert SNVs into a VCF-formatted file (software downloaded from https://github.com/liangjiaoxue/PythonNGSTools).

The CDS dataset was processed using the following steps. Filtered models CDS for fungus and endobacteria were sourced from the previously described reference genomes (*Linnemannia elongata* AD073 v1.0 (JGI Project ID: 1203123) for fungus and *Mycoavidus cysteinexigens* B1-EB (PATRIC Genome ID: 1553431.3) for endobacteria). We used BLAST to analyze the de novo assembly for CDS genes and the MUMmer package [Delcher et al., 2003] to perform variant calling on extracted CDS genes against the reference CDS genes.

Finally, the rDNA dataset was processed using the following steps. Barrnap (--kingdom euk) [Seemann, 2018] was used to identify 5S, 5.8S, 18S, and 28S subunits of rDNA from the draft fungal genomes. Then, 18S rDNA were extracted using the reference sequence (NCBI Reference Sequence: NG_070287.1). PROKKA [Seemann, 2014] was used to annotate the draft endobacteria assemblies and extract 16S rDNA. The MUMmer package [Delcher et al., 2003] was used to call fungal and endobacterial variants from the 18S and 16S rDNA, respectively.

#### Phylogenetic tree estimation

Maximum likelihood tree estimation was performed using RAxML v8.2.12 [Stamatakis, 2014] under finite-sites models of nucleotide sequence evolution. The latter consisted of the GTR [Tavaré, 1986], Jukes-Cantor Jukes and Cantor [1969], K80 [Kimura, 1980], and HKY [Hasegawa et al., 1985] models. PAUP* [Swofford, 2003] was used to conduct additional phylogenetic reconstructions using neighbor-joining (NJ) [Saitou and Nei, 1987] and the unweighted pair group method with arithmetic mean (UPGMA) algorithms [Sokal, 1958]. Multispecies coalescent model-based species tree reconstruction was performed using SVDquartet [Chifman and Kubatko, 2014]. If SVDquartet produced a tree with polytomies, the matrix rank was set to 1, 4, and 5 to produce three different tree topologies. Finally, reconstructed phylogenetic trees were midpoint rooted.

#### Cophylogenetic reconciliation and comparison of phylogenies and cophylogenies

CoRe-PA [Merkle et al., 2010] and eMPRess [Santichaivekin et al., 2021] were used to reconcile cophylogenies. Reconstructed phylogenies and cophylogenies were compared using the same calculations as in the simulation study.

### Empirical study of bobtail squids and their symbiotic bioluminescent bacteria

#### Sample acquisition and sequencing

Genomic sequence data for twenty-two samples of bobtail squids from the study of Sanchez et al. [2021] and thirty-seven *Vibrio* samples from the study of [Bongrand et al., 2020] were downloaded. Bobtail squid samples were sequenced via genome skimming to identify more than 5000 ultraconserved loci. Summary statistics for the dataset are shown in Table 8. Host-symbiont association data came from the study of Sanchez et al. [2021].

**Table 8.** Summary statistics for Bobtail squids and bioluminescent *Vibrio*.

| Organism | Data source | Summary statistics |  |  |  |  |
| --- | --- | --- | --- | --- | --- | --- |
|  |  | # taxa | Tree height | Aln length | Aln gappiness | Aln ANHD |
| Bobtail squid | Sanchez et al. [2021] | 22 | 0.1212 | 37,512 | 0.1690 | 0.0015 |
| Bioluminescent bacteria | [Bongrand et al., 2020] | 37 | 0.0109 | NA | NA | NA |

Aln: alignment, ANHD: average normalized hamming distance.

#### Reconstruction and comparison of phylogenies and cophylogenies

We reconstructed a phylogenetic tree for host taxa using the same approach as in the fungal/endobacterial dataset analysis. The bacterial symbiont phylogeny consisted of the *Vibrio* phylogeny reported by Sanchez et al. [2021]. Cophylogenetic reconciliation and comparison of estimated phylogenies and cophylogenies followed the same procedures as in the other empirical dataset analysis.

## Results

### Simulation study

#### The impact of upstream phylogenetic estimation error on downstream cophylogenetic reconciliation accuracy

Across the mixed simulation conditions, phylogenetic tree estimation returned average topological error of 7% and cophylogenetic reconstruction returned average precision of 66%. (Supplementary Figure S1 reports average topological errors of estimated species trees and cophylogenies for each model condition.) The relationship between phylogenetic and cophylogenetic estimation error was examined using linear regression: Figure 3 shows the regression models fitted to observed topological errors across replicate datasets in each model condition. The regression analyses were statistically significant in all cases (*α* = 0.05; *n* = 100), as shown in Table 9. Increasing topological error during upstream estimation was clearly associated with reduced cophylogenetic accuracy, as evidenced by consistently negative regression coefficients and average correlation coefficient of −1.96 across model conditions. We also observed varying scatter around fitted models: the coefficient of determination was highest in the mixed-gopher, mixed-stinkbug, and mixed-primate model conditions – ranging between 0.47 and 0.89 – and lower in others.

**Fig. 3.**
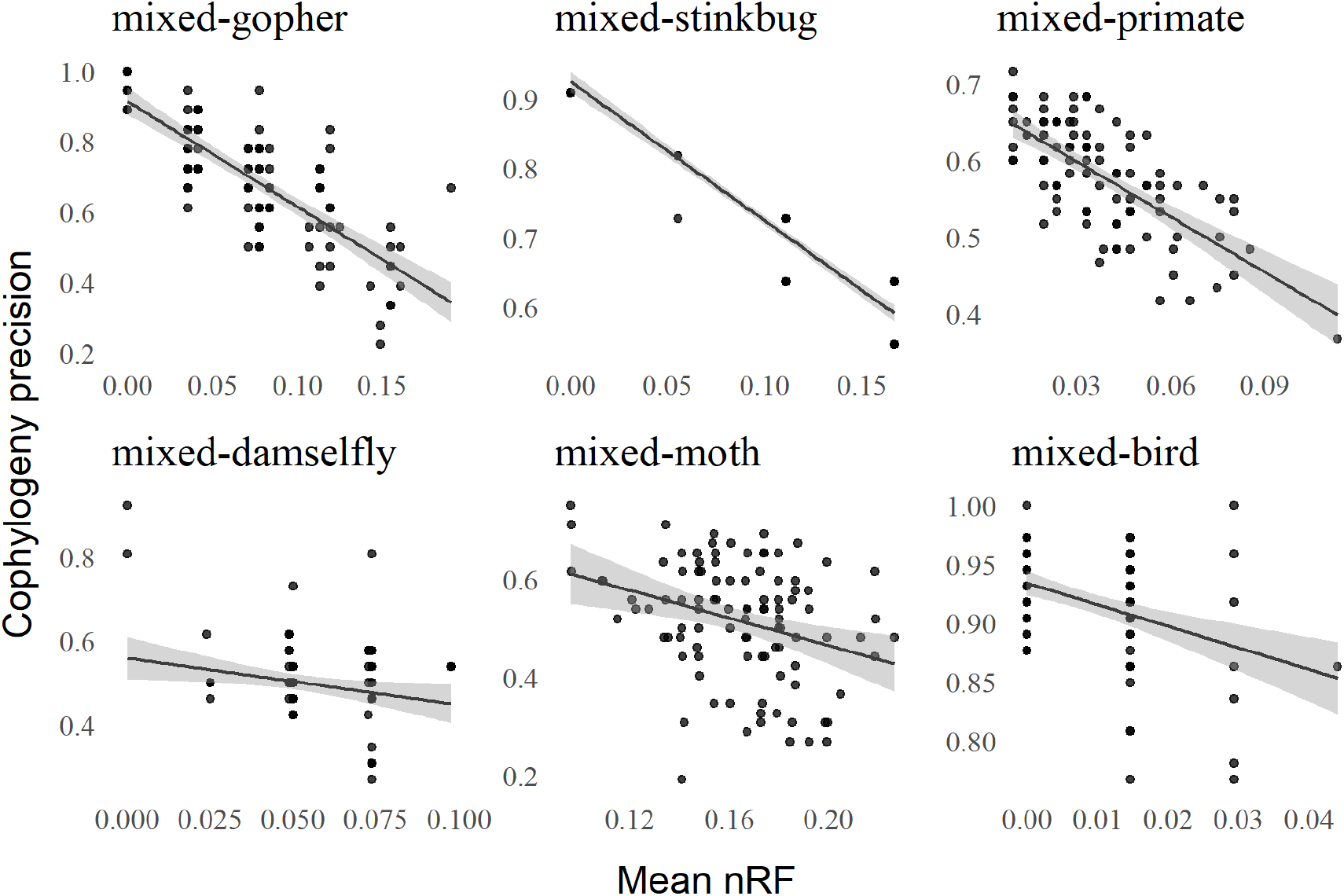
The relationship between phylogenetic and cophylogenetic estimation error on the mixed simulation conditions. For each model condition, the topological error returned by phylogenetic tree estimation (averaged across the pair of host and symbiont datasets) and the precision returned by cophylogenetic reconstruction are shown for each replicate dataset (*n* = 100). A fitted linear regression model is shown for each model condition as well, and linear regression analyses were statistically significant in all cases (*α* = 0.05; *n* = 100). The 95% confidence interval is shown in grey around the regression line.

**Table 9.** Linear regression results for mixed simulation experiments. The fitted model’s intercept (“intercept”), correlation coefficient (“B coefficient”), coefficient of determination (“R^2^”), and residual standard error (“RSE”) are shown. Statistical significance was assessed using the F-test, and uncorrected p-values (“p-value”) and corrected q-values (“q-value”) based on Benjamini-Hochberg multiple test correction [Benjamini and Hochberg, 1995] are reported (*n* = 100).

| Model conditions | Simple Linear Regression |  |  |  |  |  |
| --- | --- | --- | --- | --- | --- | --- |
|  | intercept | B coefficient | R<sup>2</sup> | RSE | p-value | q-value |
| mixed-gopher | 0.9146 | -2.9996 | 0.6406 | 0.1061 | 0.0000 | 0.0000 |
| mixed-stinkbug | 0.9254 | -2.0067 | 0.8903 | 0.0331 | 0.0000 | 0.0000 |
| mixed-primate | 0.6704 | -2.3987 | 0.4732 | 0.0511 | 0.0000 | 0.0000 |
| mixed-damselfly | 0.5590 | -1.1198 | 0.0564 | 0.0928 | 0.0173 | 0.0173 |
| mixed-moth | 0.7460 | -1.4036 | 0.1010 | 0.1146 | 0.0000 | 0.0025 |
| mixed-bird | 0.9341 | -1.8328 | 0.1663 | 0.0408 | 0.0000 | 0.0000 |

Similar outcomes were observed in the backward-time simulation experiments, as compared to the mixed simulation experiments. Upstream tree estimation returned topological error of around 10% or less (Supplementary Figure S2). Estimated cophylogeny precision was also similar – ranging around 50% to 60%. Negative and significant correlation between upstream tree error and downstream cophylogeny precision was observed on all model conditions (*α* = 0.05; *n* = 100), as shown in Figure 4. Correlation coefficients ranged between −0.644 and −0.848 (Table 10). Scatter around linear regression models was smaller than in the backward-time simulations, with coefficient of determination between 0.653 and 0.938. One minor difference between backward-time simulation experiments and mixed simulation experiments is that former the returned more consistent regression analysis results compared to the latter. We attribute the difference in part to the relative heterogeneity of the mixed simulation conditions compared to the backward-time simulation conditions.

**Fig. 4.**
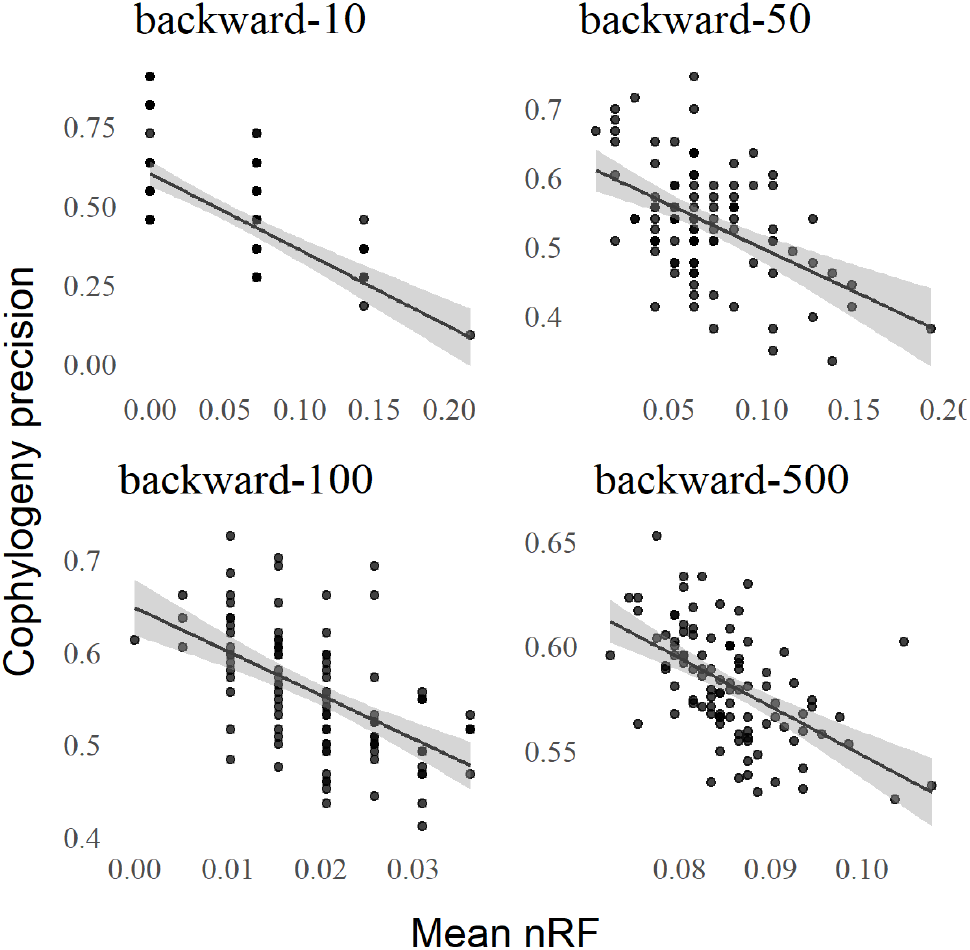
The relationship between phylogenetic and cophylogenetic estimation error on the backward-time simulation conditions. Figure layout and description are otherwise identical to Figure 3.

**Table 10.** Linear regression results for backward-time simulation experiments. Table layout and description are otherwise identical to Table 9.

| Model conditions | Simple Linear Regression |  |  |  | p-value | q-value |
| --- | --- | --- | --- | --- | --- | --- |
|  | intercept | B coefficient | R<sup>2</sup> | RSE |  |  |
| backward-10 | 0.6018 | -0.6870 | 0.6525 | 0.1644 | 0.0000 | 0.0000 |
| backward-50 | 0.6236 | -0.7010 | 0.9074 | 0.0817 | 0.0000 | 0.0000 |
| backward-100 | 0.6482 | -0.6438 | 0.9379 | 0.0545 | 0.0000 | 0.0000 |
| backward-500 | 0.7793 | -0.8475 | 0.8950 | 0.0968 | 0.0000 | 0.0000 |

Topological error of estimated phylogenies and cophylogenies varied among forward simulation conditions. The observation is due in part to heterogeneity among the empirical estimates that served as the basis for the forward-time simulation conditions. On the other hand, topological errors were somewhat higher than in the other simulation experiments: the forward-time simulation experiments returned average tree topology error of 13% and average cophylogenetic precision of 35% (Figure S3). We note that the forward-time simulation conditions do not precisely match the empirical estimates from mixed simulations, since Treeducken’s forward-time model was manually fitted. As shown in Figure 5, correlation between upstream tree estimation error and downstream cophylogeny reconstruction precision yielded similar findings as in the rest of simulation study. We observed significant and negative correlation in all forward-time simulation conditions (Table 11). Furthermore, the coefficient of determination varied across forward-time simulation conditions in a similar pattern to the mixed simulation conditions, based on shared empirical dataset estimates. The largest values were seen on forward-gopher, forward-stinkbug, and forward-primate model conditions – ranging between 0.585 and 0.744; smaller values were seen on the other model conditions.

**Fig. 5.**
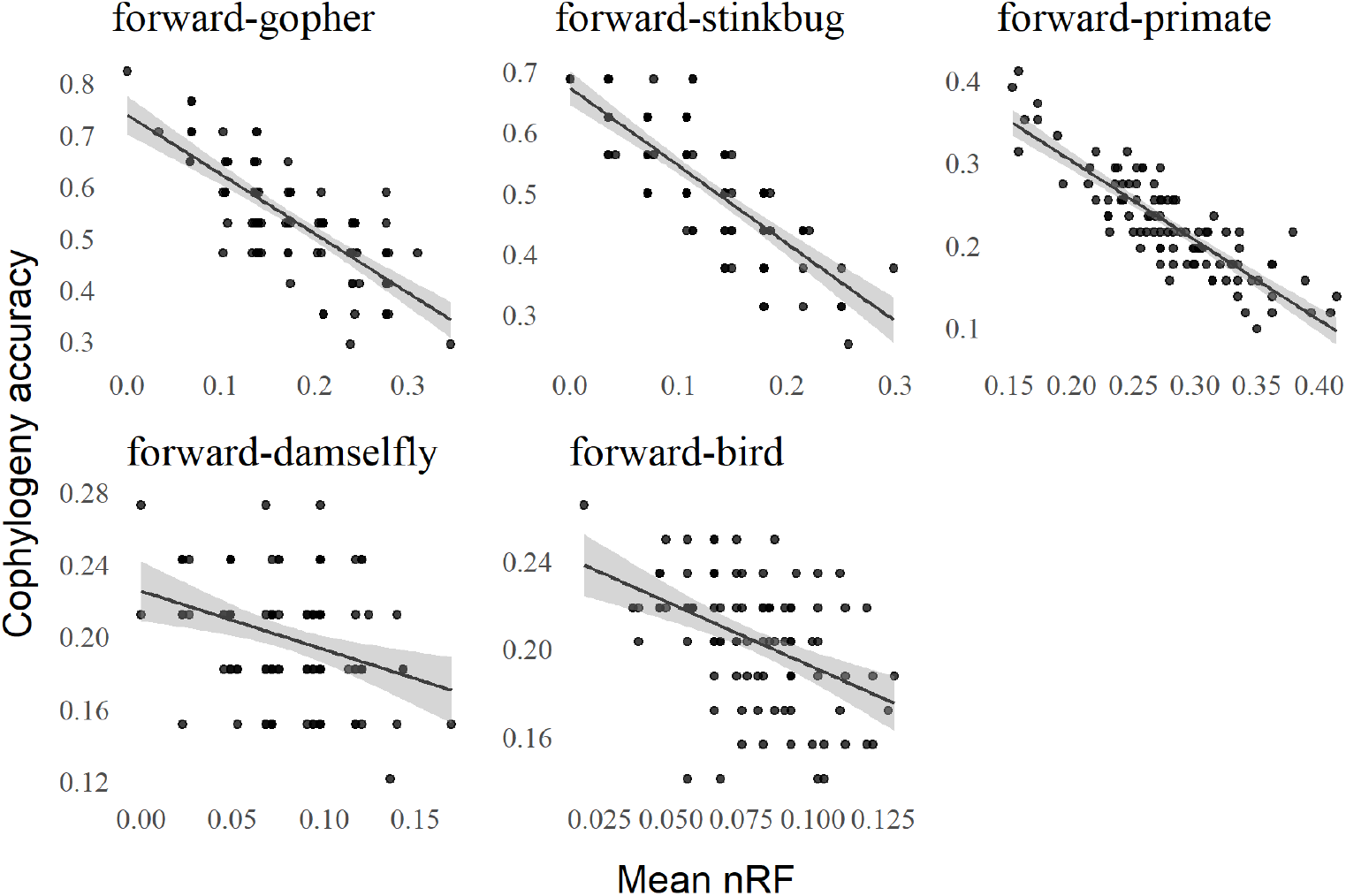
The relationship between phylogenetic and cophylogenetic estimation error on the forward-time simulation conditions. Figure layout and description are otherwise identical to Figure 3.

**Table 11.** Linear regression results for forward-time simulation experiments. Table layout and description are otherwise identical to Table 9.

| Model conditions | Simple Linear Regression |  |  |  |  |  |
| --- | --- | --- | --- | --- | --- | --- |
|  | intercept | B coefficient | R<sup>2</sup> | RSE | p-value | q-value |
| forward-gopher | 0.7385 | -1.1485 | 0.5854 | 0.0680 | 0.0000 | 0.0000 |
| forward-stinkbug | 0.6729 | -1.2848 | 0.6171 | 0.0632 | 0.0000 | 0.0000 |
| forward-primate | 0.4968 | -0.9702 | 0.7442 | 0.0312 | 0.0000 | 0.0000 |
| forward-damselfly | 0.2252 | -0.3232 | 0.1035 | 0.0326 | 0.0011 | 0.0011 |
| forward-bird | 0.2495 | -0.5780 | 0.1129 | 0.0141 | 0.0000 | 0.0000 |

#### Impact of evolutionary divergence on the relationship between phylogenetic and cophylogenetic reconstruction accuracy

For each set of backward-time and forward-time simulation conditions (Figure 6 and Figure 7 respectively), we found that phylogenetic and cophylogenetic estimation error was negatively and significantly correlated as the tree height parameter *h* varied between 0.1 and 10. Regression analysis returned correlation coefficients between −0.899 and −0.220, and coefficients of determination between 0.957 and 0.169 (Tables 12 and 13). Both upstream and downstream topological error was lowest for the smallest *h* settings (i.e., 0.1, 0.5, and 1.0). As the height *h* increased, both topological errors increased in tandem, and both were largest on simulations with height *h* = 10. The latter was likely at saturation, as topological errors tended to be maximal. Similar outcomes were observed in the corresponding mixed simulation experiments with varying tree height *h*, as shown in Figure 8 with regression analysis results listed in Table 14. The effect of increasing *h* on topological error was more complicated and non-linear in some cases. This was in part due to heterogeneity of empirical estimates used for parametric resampling, unlike the fully *in silico* simulations used elsewhere in the simulation study.

**Fig. 6.**
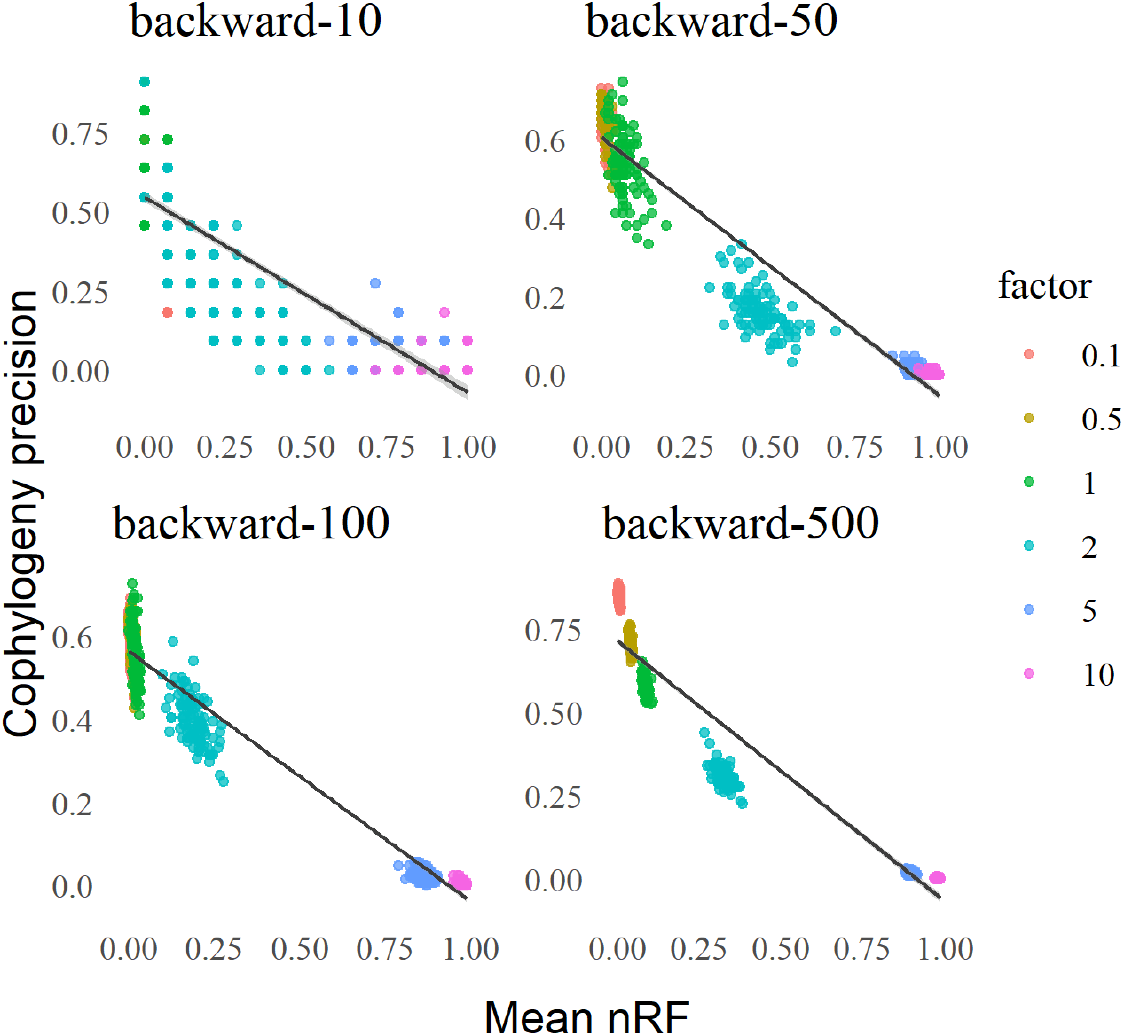
Backward-time simulation experiments: the impact of evolutionary divergence on phylogenetic and cophylogenetic estimation error. Figure layout and description are otherwise identical to Figure 8.

**Fig. 7.**
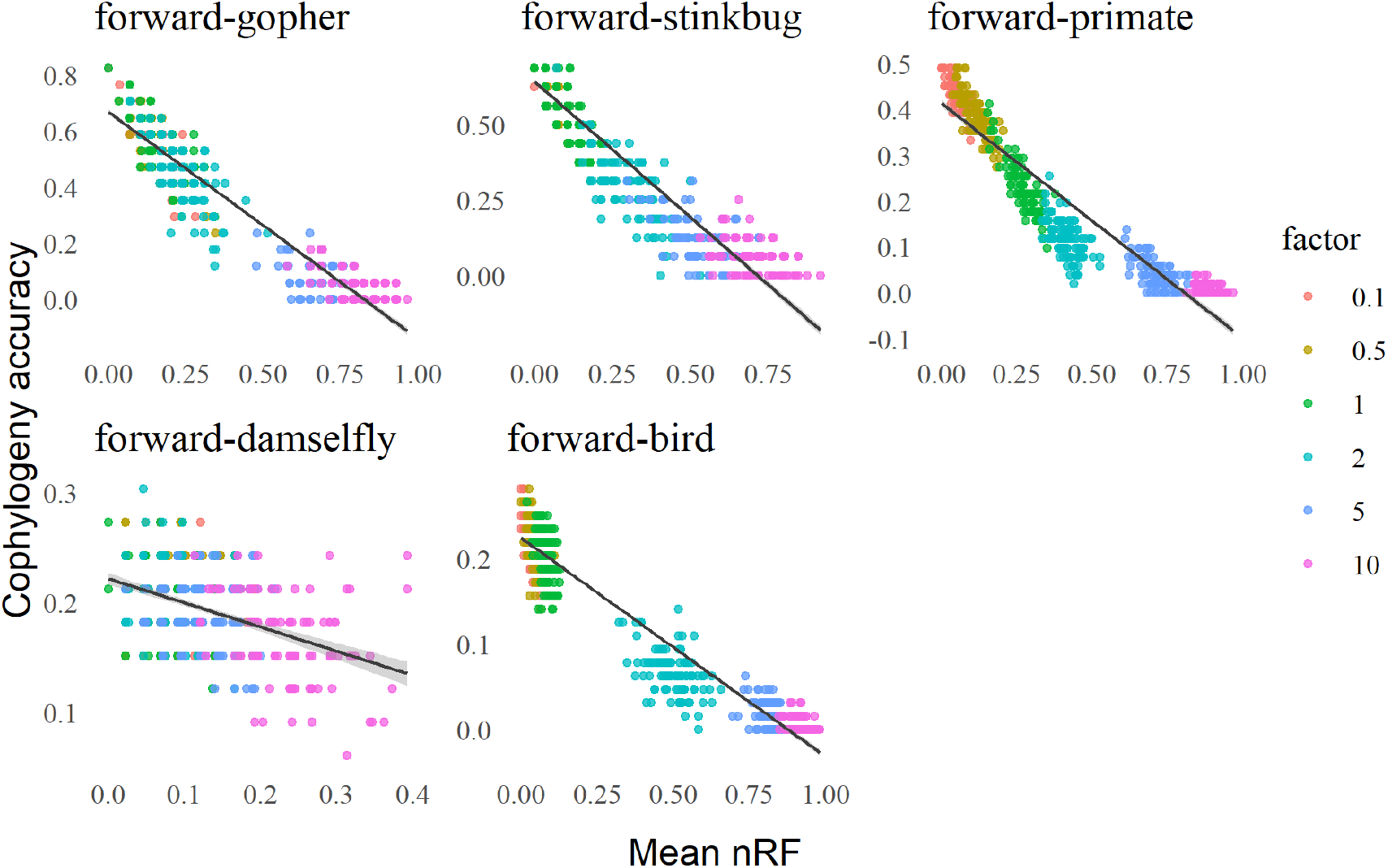
Forward-time simulation experiments: the impact of evolutionary divergence on phylogenetic and cophylogenetic estimation error. Figure layout and description are otherwise identical to Figure 8.

**Fig. 8.**
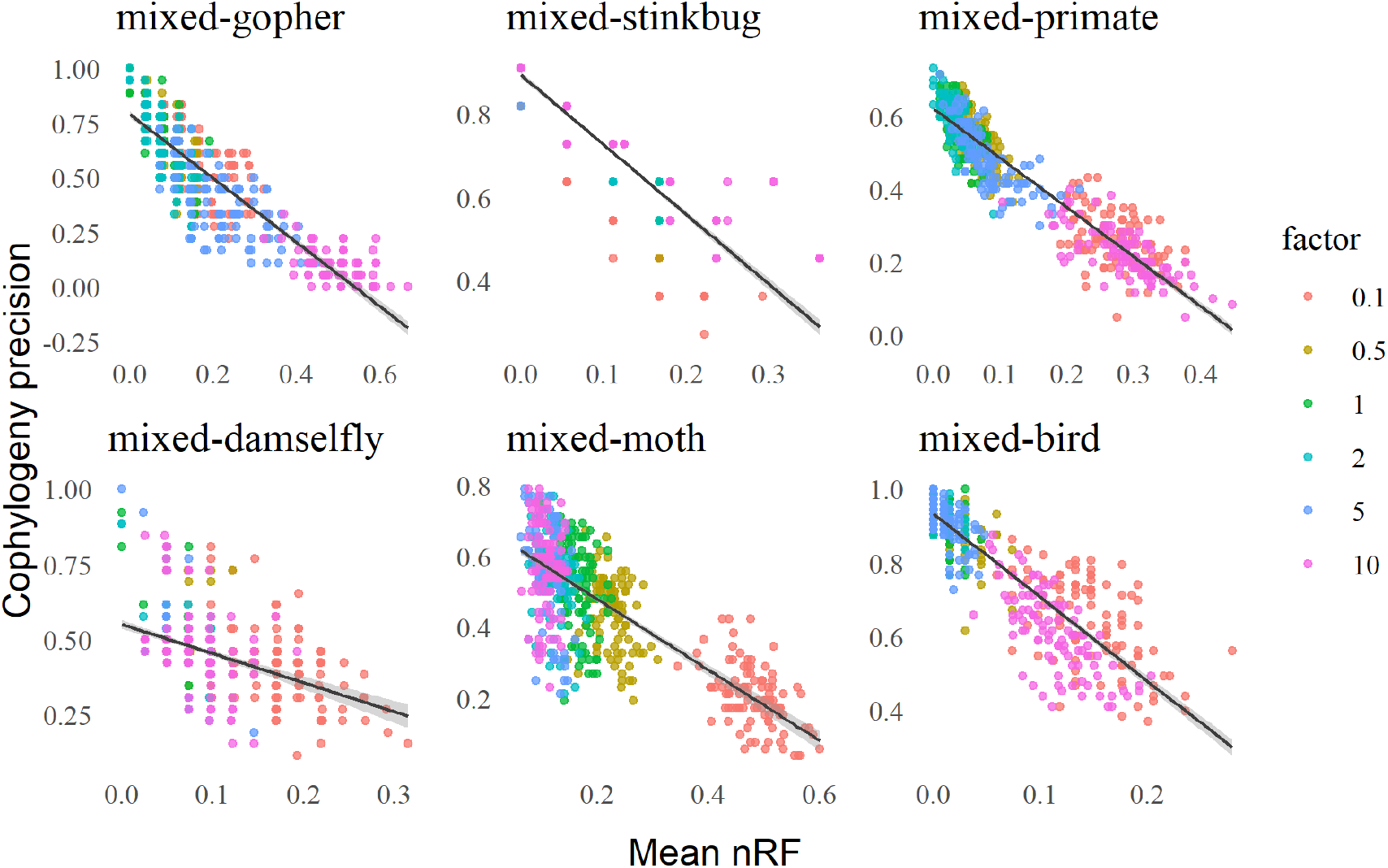
Mixed simulation experiments: the impact of evolutionary divergence on phylogenetic and cophylogenetic estimation error. Estimation error was assessed based upon average topological error of estimated trees (averaged across the pair of host and symbiont datasets) and cophylogenetic precision. Model tree branch lengths were scaled by height parameter *h* (“factor”); data points for a given setting of *h* are distinguished by a distinct color. A fitted linear regression model is shown for each mixed simulation condition (*n* = 600).

**Table 12.**
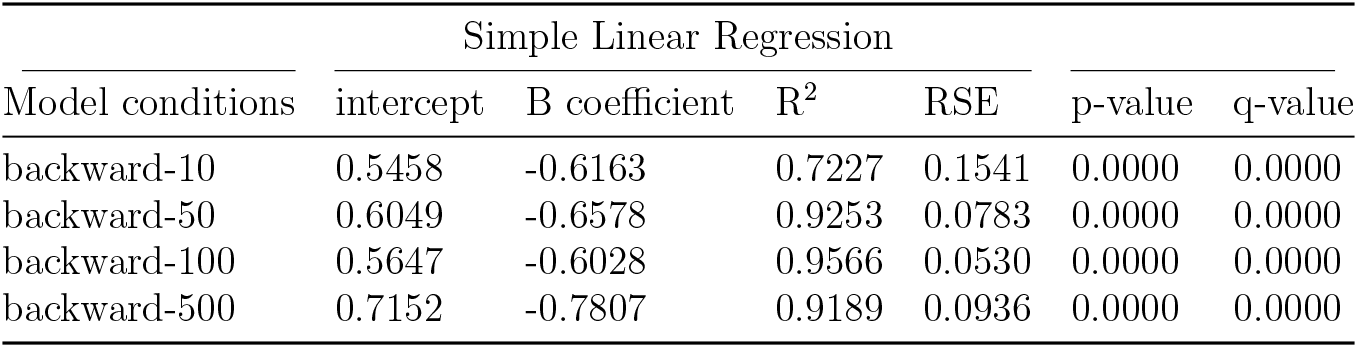
Linear regression results for evolutionary divergence, backward-time simulation experiments. Table layout and description are otherwise identical to Table 9.

**Table 13.** Linear regression results for evolutionary divergence, forward-time simulation experiments. Table layout and description are otherwise identical to Table 9.

| Model conditions | Simple Linear Regression |  |  |  | p-value | q-value |
| --- | --- | --- | --- | --- | --- | --- |
|  | intercept | B coefficient | R<sup>2</sup> | RSE |  |  |
| forward-gopher | 0.6677 | -0.8078 | 0.9091 | 0.0738 | 0.0000 | 0.0000 |
| forward-stinkbug | 0.6429 | -0.8991 | 0.9091 | 0.0777 | 0.0000 | 0.0000 |
| forward-primate | 0.4133 | -0.5121 | 0.8796 | 0.0584 | 0.0000 | 0.0000 |
| forward-damselfly | 0.2217 | -0.2200 | 0.1693 | 0.0344 | 0.0000 | 0.0000 |
| forward-bird | 0.2241 | -0.2553 | 0.9317 | 0.0257 | 0.0000 | 0.0000 |

**Table 14.**
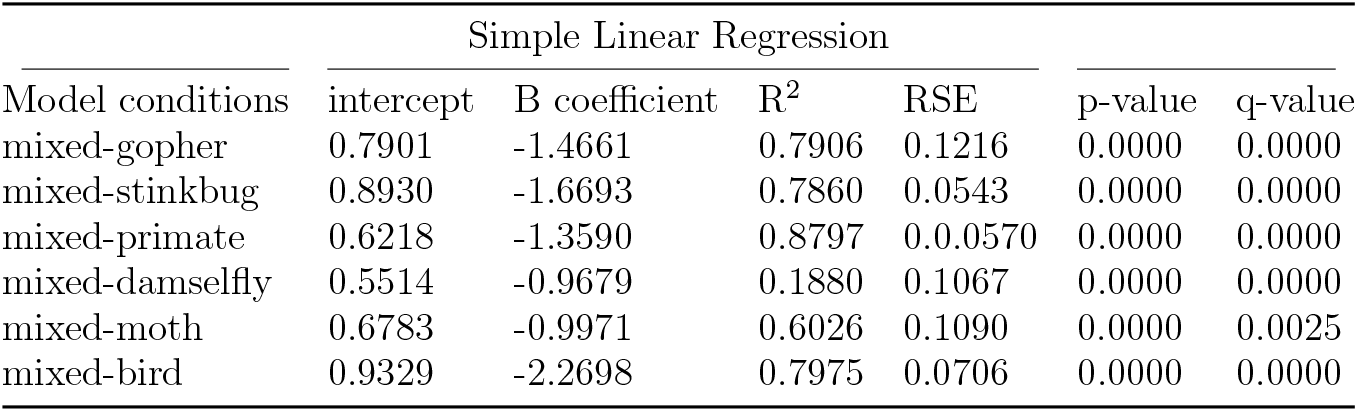
Linear regression results for evolutionary divergence, mixed simulation experiments. Table layout and description are otherwise identical to Table 9.

### Empirical study

#### Soil-associated fungi and their bacterial endosymbionts

Topological disagreements among estimated phylogenies were higher than in the simulation study (Supplementary Figure S4); a similar outcome was observed among estimated cophylogenies. This is by design: the empirical study utilized a wide array of phylogenetic reconstruction methods with varying estimation accuracy. The design choice provides an indirect means to vary the topological accuracy of input phylogenies and then observe its effects on downstream cophylogenetic estimation, in contrast to the direct control and ground truth enabled by *in silico* simulations. We analyzed the relationship between phylogenetic and cophylogenetic estimation error using linear regression (Figure 9). Consistent with the simulation study, we observed that greater topological agreement in the former set of inputs was significantly associated with greater topological agreement of the latter output (*α* = 0.05; *n* = 114, *n* = 137, and *n* = 78 for the full-assembly, CDS, and rDNA datasets, respectively). The full assembly dataset analysis returned a regression coefficient of −2.067 and coefficient of determination of 0.678, which is also in line with the simulation study (Table 15). Similar outcomes were observed on the smaller CDS and rDNA datasets.

**Fig. 9.**
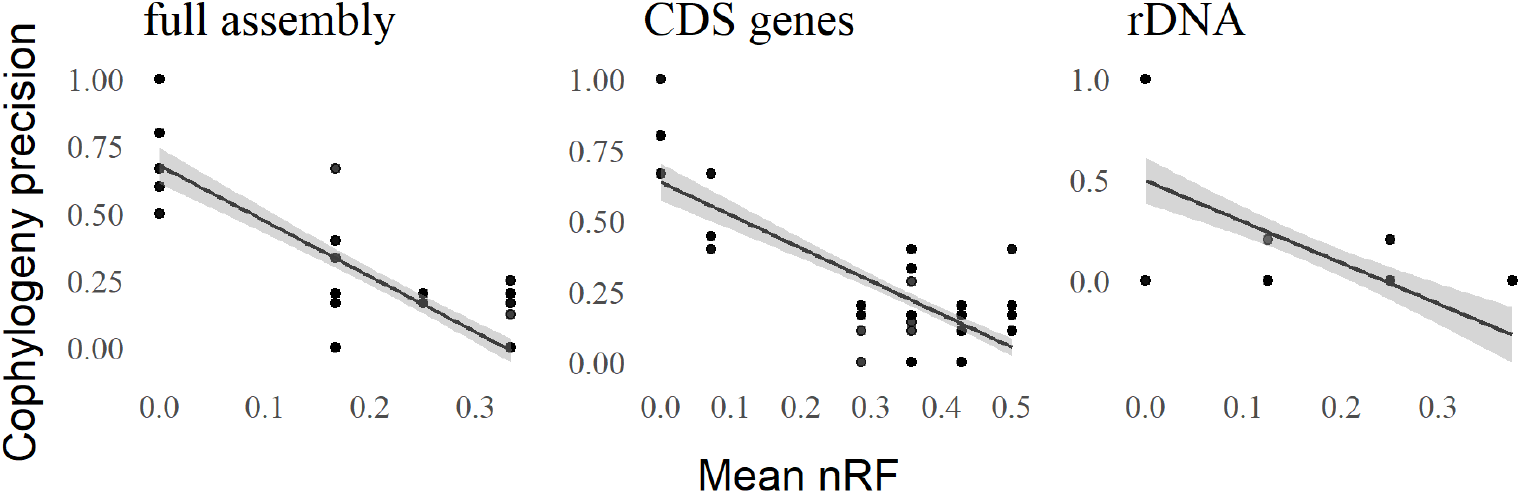
Topological discordance among phylogenetic and cophylogenetic estimates for soil-associated fungi and their bacterial endosymbionts. A range of different methods were used to estimate phylogenetic trees for host taxa, and similarly for symbiont taxa; for a given set of taxa (either host or symbiont), pairwise topological discordance among the resulting tree estimates was assessed based on normalized Robinson-Foulds distance. Then, a cophylogeny was reconstructed using each pair of host/symbiont trees that was estimated using a given phylogenetic tree estimation method (along with the known host/symbiont associations); each pair of estimated cophylogenies was compared based on cophylogenetic precision. A scatterplot and fitted linear regression model is shown for the full-assembly, CDS, and rDNA datasets (*n* = 114, *n* = 137, and *n* = 78, respectively, where CoRe-PA returned multiple estimates in the event of co-optimal solutions).

**Table 15.** Linear regression results for soil-associated fungi and their bacterial endosymbionts. As noted in Figure 9, linear regression was used to analyze the agreement between phylogenetic and cophylogenetic estimates, where the former varied due to the choice of phylogenetic estimation method used and the latter’s input was based on the former. Results for linear regression analyses are reported in a manner and layout identical to those in Table 9.

| VCF Datasets | Simple Linear Regression |  |  |  | p-value | q-value |
| --- | --- | --- | --- | --- | --- | --- |
|  | intercept | B coefficient | R<sup>2</sup> | RSE |  |  |
| full assembly | 0.6781 | -2.0672 | 0.6723 | 0.1740 | 0.0000 | 0.0000 |
| CDS genes | 0.6370 | -1.1656 | 0.5839 | 0.1314 | 0.0000 | 0.0000 |
| rDNA | 0.4954 | -2.0426 | 0.3919 | 0.2841 | 0.0000 | 0.0000 |

#### Bobtail squids and their symbiotic bioluminescent bacteria

Topological disagreements among species cophylogenies and resulting cophylogenetic reconciliations were somewhat smaller than those observed on fungal/endosymbiont dataset (Supplementary Figure S5). Another key difference concerns host/symbiont associations: relatively few squid hosts were associated with most bacterial symbionts. Still, we observed a similar relationship between upstream phylogenetic estimation agreement and downstream cophylogeny precision (Figure 10). Linear regression analyses returned significant and negative correlation (*α* = 0.05; *n* = 100), with correlation coefficient of −0.449, intercept of 0.841, F-test p-value < 10^-12^, coefficient of determination of 0.213, and residual standard error of 0.109.

**Fig. 10.**
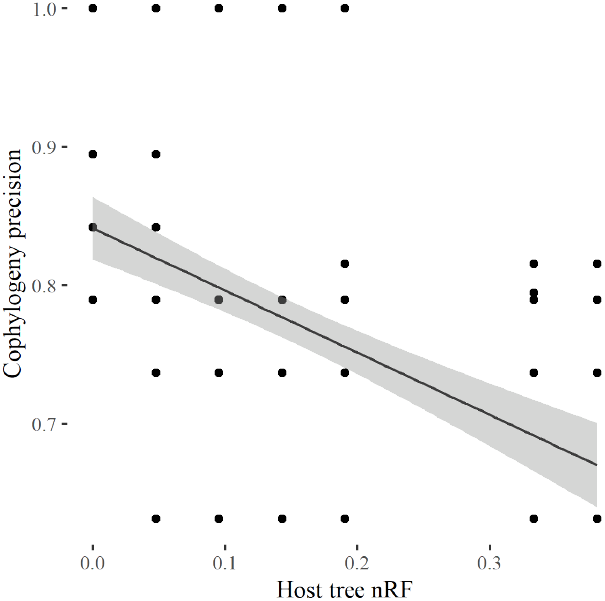
Topological discordance among phylogenetic and cophylogenetic estimates for bobtail squids and their bioluminescent symbionts. Figure description and layout are otherwise identical to Figure 9.

## Discussion

Across all forward-time simulation experiments, correlation between upstream phylogenetic estimation error and downstream cophylogenetic estimation accuracy was significant and consistently negative. As the former increased, the latter would degrade. The mixed and backward-time simulation experiments and empirical dataset analyses also returned a consistent outcome: namely, a significant and negatively correlated relationship between upstream phylogenetic reconstruction error and downstream cophylogenetic estimation reproducibility. Furthermore, the expanded simulation study experiments that focused on varying evolutionary divergence (while fixing other experimental factors) refined our study’s primary finding. We found that evolutionary divergence plays a key role in modulating upstream and downstream estimation error in tandem. Of course, other factors also play a role (e.g., taxon sampling, coevolutionary event distribution, evolutionary and coevolutionary model mis-specification, etc.), and the relationship between phylogenetic and cophylogenetic reconstruction is quite complex. Heterogeneity among simulation conditions due to these factors helps to explain some of the more minor differences among experimental outcomes. Nevertheless, our primary finding – that phylogenetic estimation error strongly impacts downstream cophylogenetic reconciliation accuracy – was robust to these factors.

We note that the event-based cophylogeny reconstruction methods under study by default assign the lowest cost penalty to cospeciation events, which has been theorized to bias these software towards cospeciation [Nuismer and Week, 2019, Vienne et al., 2013]. The forward-time simulation experiment revealed that this potential bias has consequences. The forward-bird and forward-damselfly model condition included a lower proportion of cospeciation events compared to other forward-time simulation conditions. On these model conditions, we observed cophylogenetic reconciliation accuracy of at most 28% and 27%, respectively, which were the lowest in the forward-time simulation experiments. In contrast, the forward-gopher and forward-stinkbug simulation experiments yielded cophylogenetic reconstruction precision of at most 82% and 69%, respectively. The comparison underscores the complexity of the cophylogeny reconstruction problem.

We note a key difference between the simulation study and the empirical study. A primary advantage of the former is the ability to benchmark against ground truth. But the latter is inherently more complex and nuanced than the former. For example, the two systems in our empirical study are models sampled along a continuum of symbiotic coevolution modes: from open – as in the case of bobtail squids and their bioluminescent symbionts [Perreau and Moran, 2022] – to mixed to closed – as in the case of early diverging fungi and their endosymbionts [Pawlowska et al., 2018]. Depending on the taxa under study, it is plausible that symbiotic coevolution may switch between different modes along a phylogeny (e.g., from closed to mixed). But we are not aware of any suitable non-homogeneous cophylogenetic models and we also lack a basic understanding of their theoretical properties (e.g., statistical identifiability). The gap between natural symbiotic coevolution and emerging statistical cophylogenetic models represent an immediate opportunity for advanced model development.

## Conclusion

This study demonstrated the major effect that phylogenetic estimation error has on downstream cophylogenetic reconstruction accuracy. The finding was consistently observed throughout the simulation study experiments. Empirical analyses of two genomic sequence datasets for models of symbiosis also revealed that variable phylogenetic tree estimation quality decreased reproducibility of cophylogenetic estimation.

We conclude with thoughts on future research directions. In addition to the previous discussion about future cophylogenetic modeling efforts, our study points to another urgent necessity. New cophylogeny reconstruction methods that explicitly account for input species tree topological error are needed to address the core issue in our study. Statistical methods that reconstruct a cophylogeny using an input species tree distribution or simultaneously co-estimate species trees and a cophylogeny would be ideal. But an important prerequisite must be addressed first – realistic models of coevolution (as discussed above) that also permit tractable statistical calculations. And statistical efficiency of inference and learning algorithms under the new models is also paramount. As noted above, there have been some past research efforts in this direction (e.g., Baudet et al. [2015]’s non-rate-based statistical formulation of the Duplication-Transfer-Loss model); more recently, Treeducken’s forward-time model [Dismukes and Heath, 2021] is a new and promising coalescent-based alternative to existing models. However, we anticipate that computational tractability (even using approximate inference techniques like approximate Bayesian computation, pseudolikelihood maximization, or others) will be a truly formidable challenge. As a temporary workaround, we propose that researchers adopt more intensive species tree reconstruction as best practices in a cophylogenetic study. For example, we recommend that researchers select more intensive local optimization heuristic settings for addressing the computationally difficult tree reconstruction problems in this study and in the state of the art. Where available, more high-quality biomolecular sequence data can also help, assuming that suitable methods can be used to account for the complex interplay of evolutionary processes – substitutions, sequence insertion and deletion, genetic drift and incomplete lineage sorting, and more – that arises in this setting.

## Data Availability

Updated versions of the study data and software scripts underlying this article are available in the public GitLab repository at https://gitlab.msu.edu/liulab/cophylogeny-species-tree-quality-performance-study-data-scripts. An archival snapshot of the study data and software scripts has been uploaded to Figshare and can be accessed at https://doi.org/10.6084/m9.figshare.21713996.v1.

## Acknowledgment

This research has been supported in part by the National Science Foundation (2144121, 2214038, 1740874, 1737898 to KJL) and MSU (EEB summer fellowship to JZ). All computational experiments and analyses were performed on the MSU High Performance Computing Center, which is part of the MSU Institute for Cyber-Enabled Research.

