## Supplementary Online Materials for "The Impact of Species Tree Estimation Error on Cophylogenetic Reconstruction"

### Supplementary document

**S5 Commands to run cophylogenetic reconciliation software** **12**

**S6 Commands used in empirical experiments** **12**

**S7 Commands used in simulation experiments** **14**

**S8 Commands to run simulator software** **14**

**S9 Custom-modified Treeducken code** **15**

#### S1 Bar plots

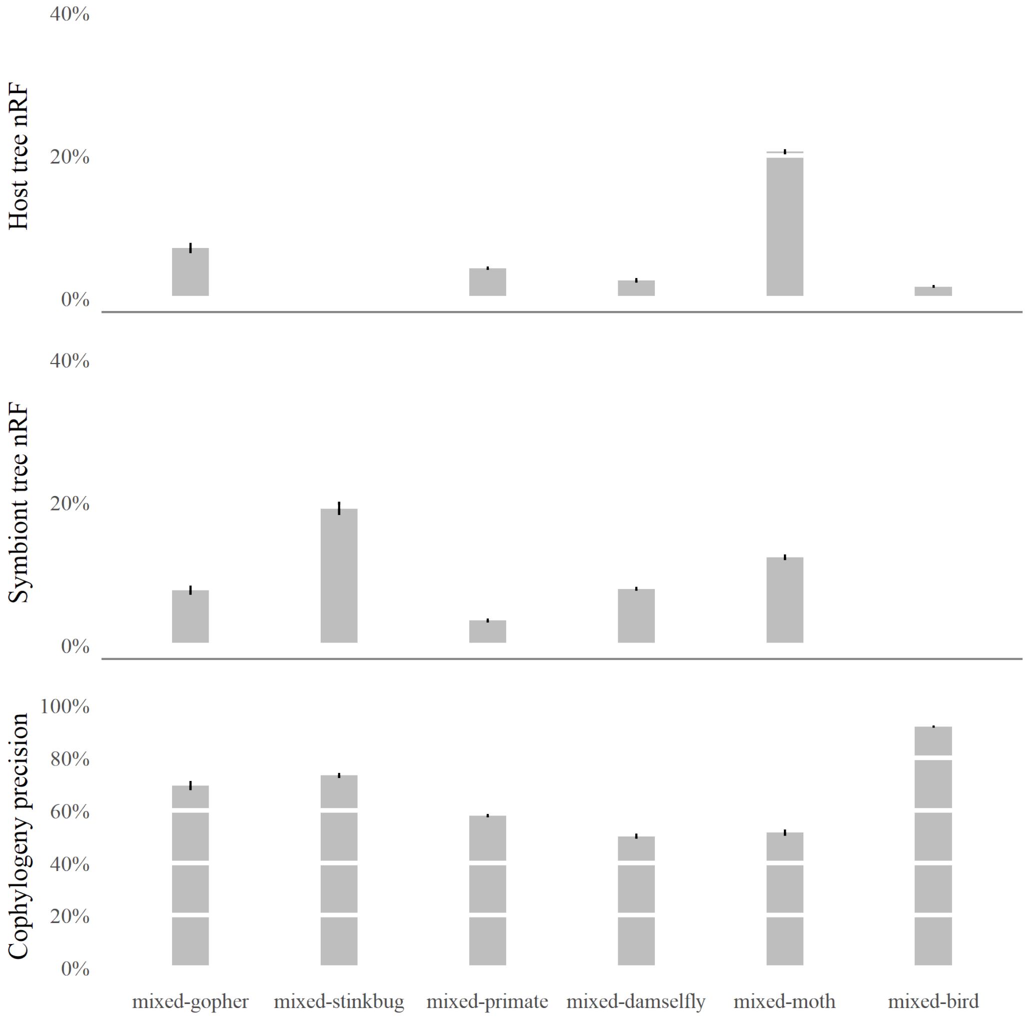

Figure S1: **For each mixed simulation condition, host tree topology error, average symbiont tree topology error, and cophylogenetic precision are shown.** Averages are reported across all experimental replicate for each model condition ($n = 100$). Standard error bars are shown.

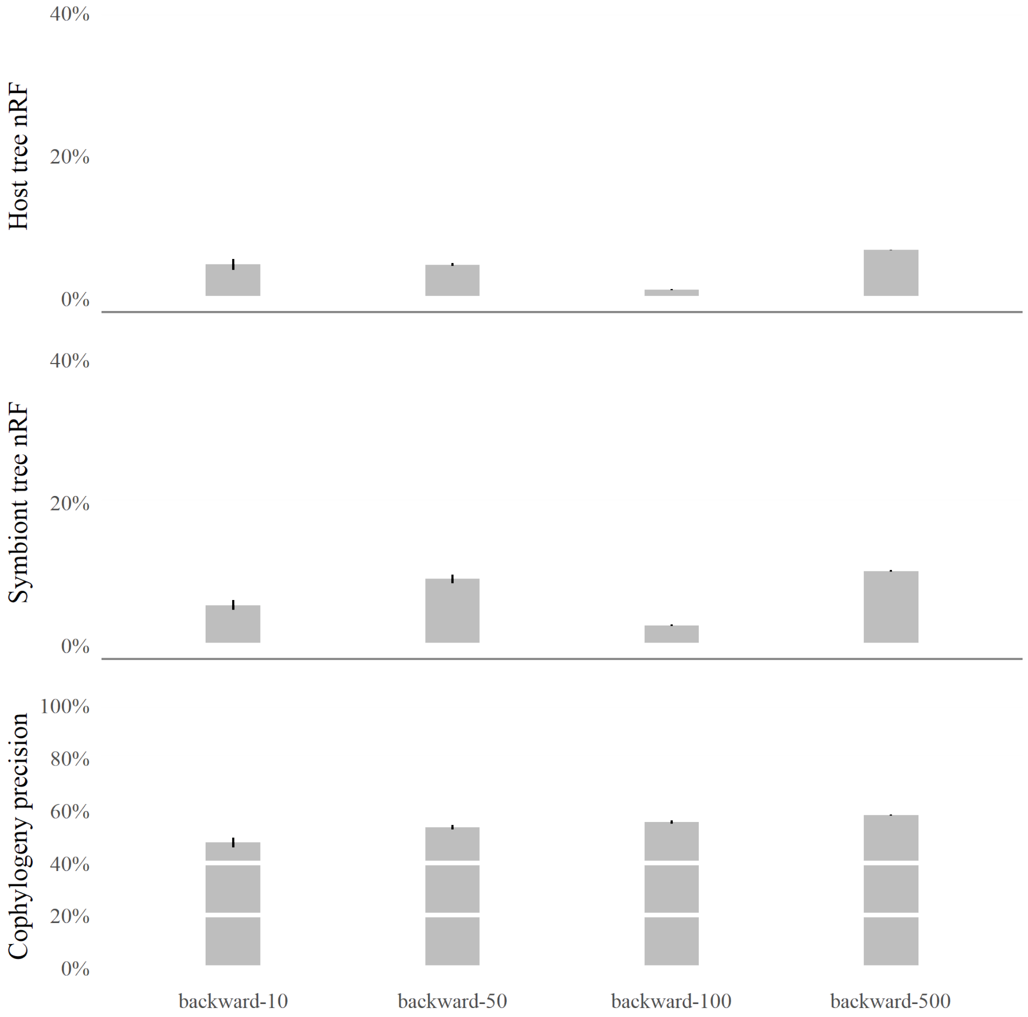

Figure S2: **Backward simulation bar graphs for average host tree topology error, average symbiont tree topology error, and average cophylogenetic precision.** Averages are reported across all experimental replicate for each model condition ($n = 100$). Error bars visualize standard error.

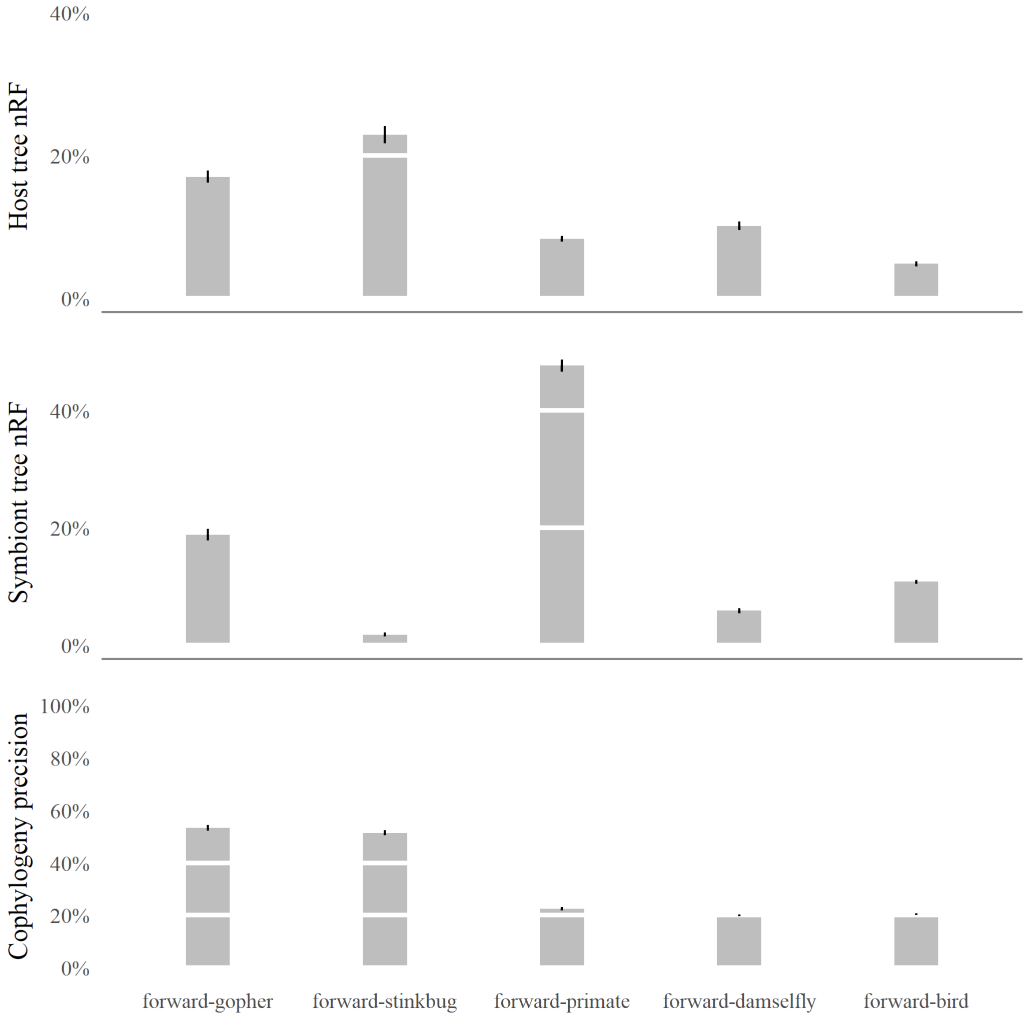

Figure S3: **Forward simulation bar graphs for average host tree topology error, average symbiont tree topology error, and average cophylogenetic precision.** Error bars visualize standard error. Averages are reported across all experimental replicate for each model condition ($n = 100$).

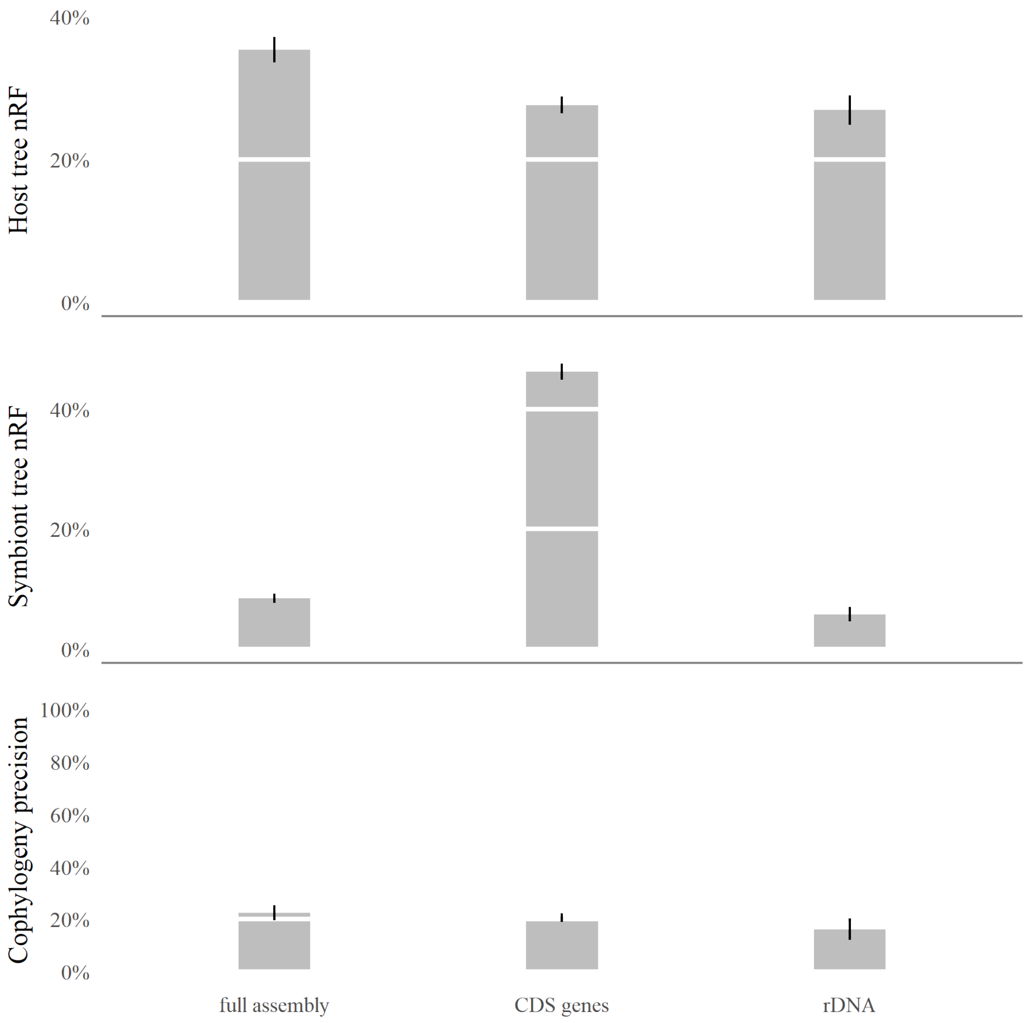

Figure S4: **Bar graphs for *Mortierella spp.* and endobacteria dataset.** Top to bottom: Average host tree error, average symbiont tree error, and average cophylogenetic precision ($n = 100$).

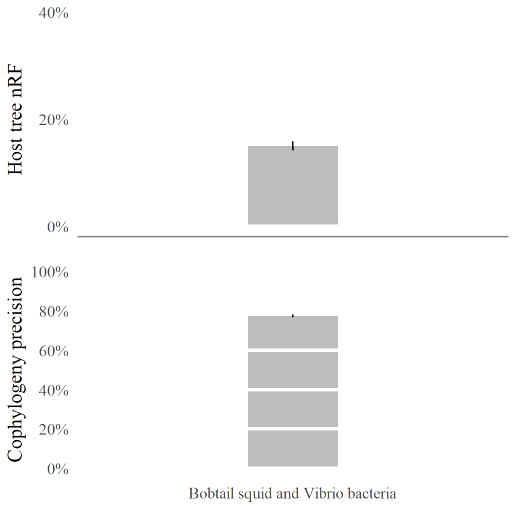

Figure S5: **Bar plots for bobtail squid and *Vibrio* dataset for average host tree error and average cophylogenetic precision.** Averages are reported across all experimental replicates ($n = 100$).

#### S2 Comparison between default event cost penalty and alternative event penalties

**Experiments on event costs used for co-phylogenetic reconciliation.** Reconciliations were assessed with different event costs estimated by COALA and CoRe-PA. On all forward-time simulation model, we found that the alternative event costs did not outperform the default event costs used by eMPRess (Figure S7). A similar outcome was observed on the mixed simulation conditions (Figure S6). For this reason, our performance study primarily utilizes default event to perform eMPRess analyses.

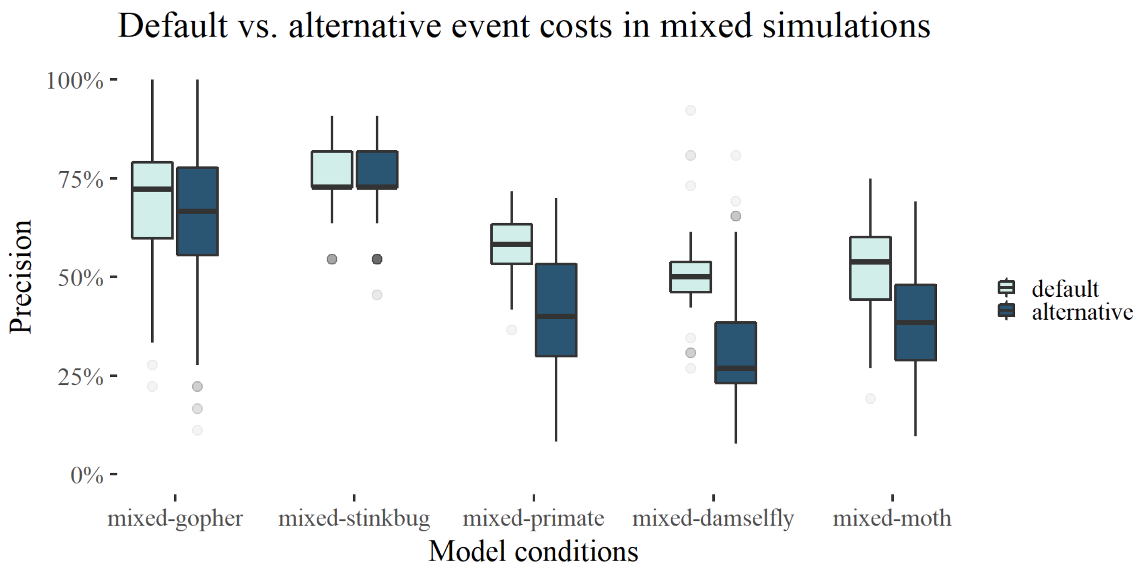

Figure S6: **Effect of using default event cost versus COALA and CoRe-PA-estimated event frequencies in eMPRess reconciliations for mixed simulations.** Co-phylogenetic precision is reported across all model condition, each with $n = 100$ experimental replicates.

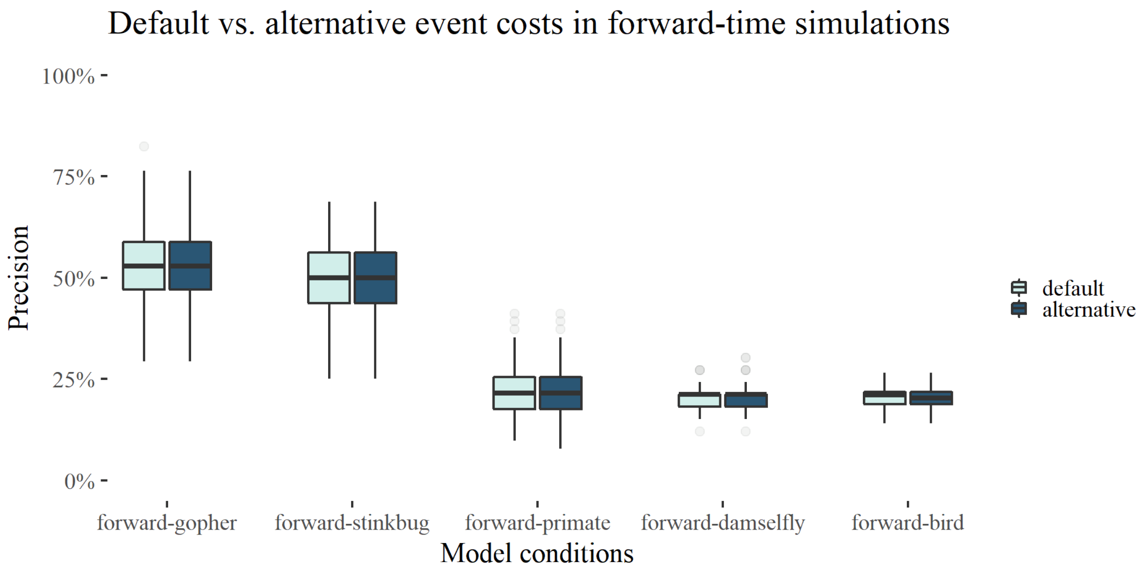

Figure S7: **Effect of using default event cost versus COALA and CoRe-PA-estimated event frequencies in eMPRess reconciliations for forward simulations.** Co-phylogenetic accuracy is reported across all model condition, each with $n = 100$ experimental replicates.

### S3 Additional empirical study experiments

#### S3.1 *Mortierella spp.* and endosymbiont: unpruned datasets

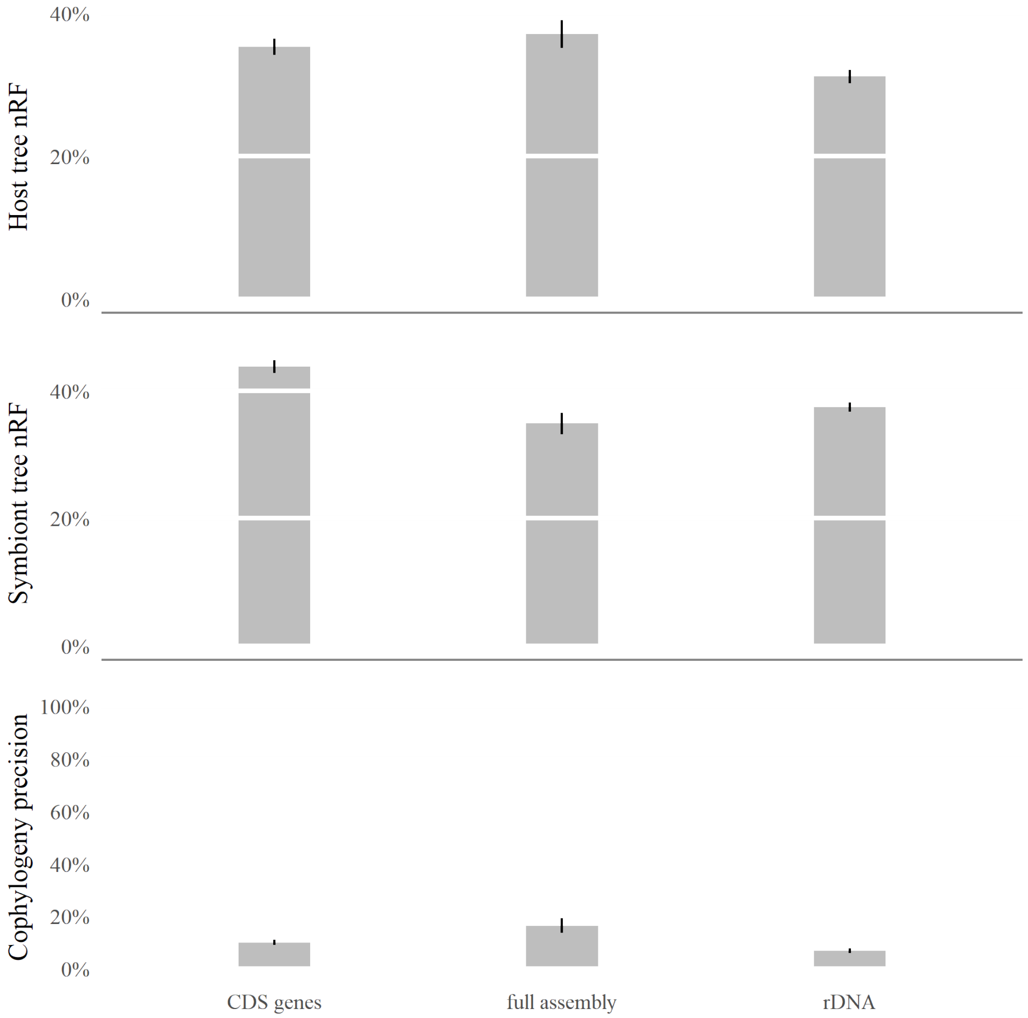

Figure S8: **Bar graphs for unpruned *Mortierella spp.* and endobacteria datasets.** Top to bottom: Average host tree error, average symbiont tree error, and average cophylogenetic precision ($n = 100$).

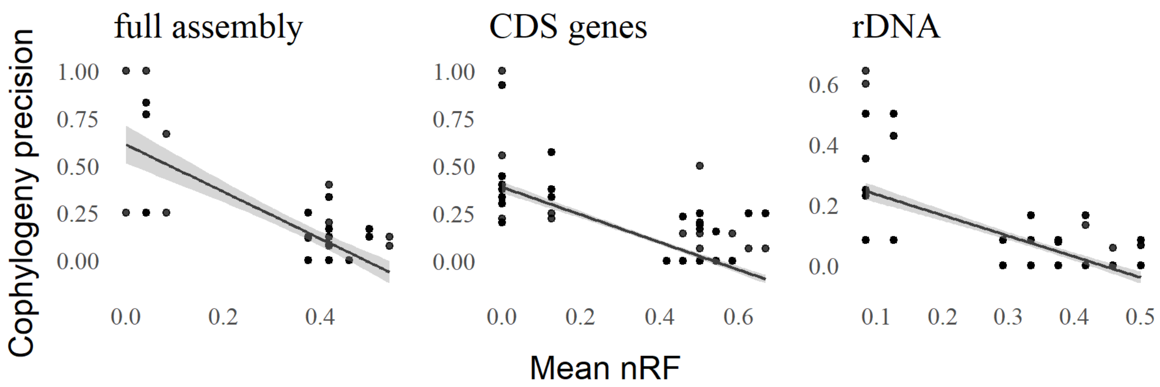

Figure S9: **Topological discordance among phylogenetic and co-phylogenetic estimates for soil-associated fungi and their bacterial endosymbionts: unpruned VCF datasets.** A scatterplot and fitted linear regression model is shown for the unpruned full-assembly, CDS, and rDNA datasets ($n = 76$, $n = 321$, and $n = 251$, respectively).

| | Simple Linear Regression | | | | | |
|---|---|---|---|---|---|---|
| VCF Datasets | intercept | B coefficient | $R^2$ | RSE | p-value | q-value |
| full assembly | 0.6104 | -1.2402 | 0.5546 | 0.1616 | 0.0000 | 0.0000 |
| CDS genes | 0.3879 | -0.7265 | 0.5706 | 0.1139 | 0.0000 | 0.0000 |
| rDNA | 0.3029 | -0.6841 | 0.4372 | 0.0986 | 0.0000 | 0.0000 |

Table S1: **Linear regression results for soil-associated fungi and their bacterial endosymbionts: unpruned datasets.** Linear regression was used to analyze the agreement between phylogenetic and co-phylogenetic estimates, where the former varied due to the choice of phylogenetic estimation method used and the latter's input was based on the former.

#### S3.2 *Mortierella spp.* and endosymbiont: bootstrap experiment

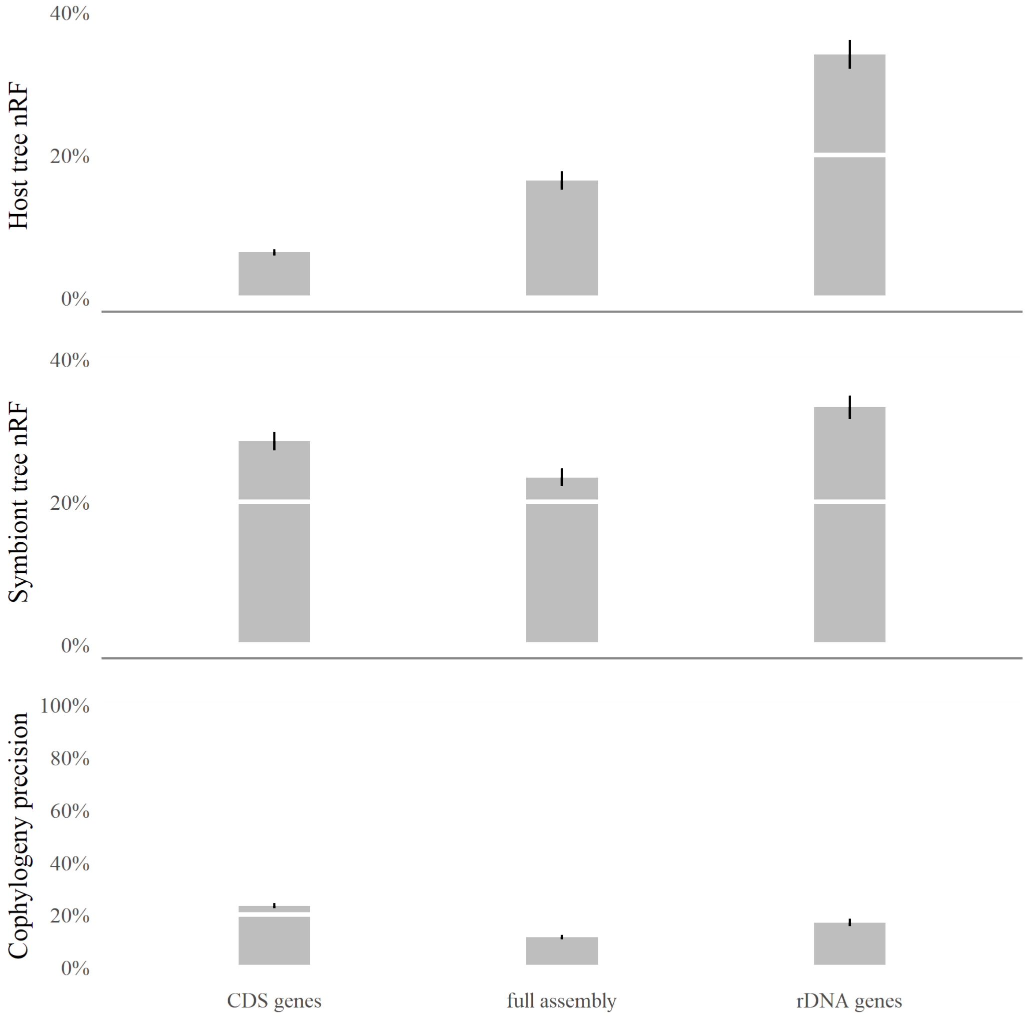

Figure S10: **Bar graphs for *Mortierella spp.* and endobacteria datasets in bootstrap experiment.** Top to bottom: Average host tree error, average symbiont tree error, and average cophylogenetic precision ($n = 100$).

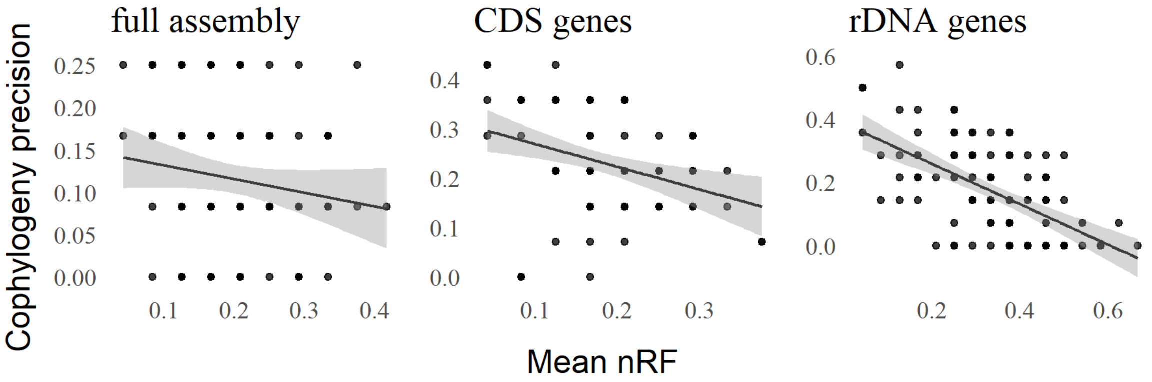

Figure S11: **Topological discordance among phylogenetic and co-phylogenetic estimates for soil-associated fungi and their bacterial endosymbionts in bootstrap experiment.** A scatterplot and fitted linear regression model is shown for the unpruned full-assembly, CDS, and rDNA datasets ($n = 76$, $n = 321$, and $n = 251$, respectively).

| | Simple Linear Regression | | | | | |
|---|---|---|---|---|---|---|
| VCF Datasets | intercept | B coefficient | $R^2$ | RSE | p-value | q-value |
| Bootstrap full assembly | 0.1477 | -0.1598 | 0.0242 | 0.0847 | 0.1223 | 0.1223 |
| Bootstrap CDS genes | 0.3151 | -0.4597 | 0.0963 | 0.0986 | 0.0017 | 0.0033 |
| Bootstrap rDNA | 0.3854 | -0.6377 | 0.3586 | 0.1164 | 0.0000 | 0.0000 |

Table S2: **Linear regression results for soil-associated fungi and their bacterial endosymbionts: bootstrap experiment.** Linear regression was used to analyze the agreement between phylogenetic and co-phylogenetic estimates, where the former varied due to the choice of phylogenetic estimation method used and the latter's input was based on the former.

### S4 Experiments with CoRe-PA

Following the simulation methods section in the main paper, we reproduced the same experimental conditions and reconstructed the cophylogenies using CoRe-PA [Merkle et al., 2010] instead of eMPRess. In general, we obtained similar findings in CoRe-PA experiments as in the eMPRess experiments, thus confirming our findings in the main manuscript.

#### S4.1 Mixed simulation results with CoRe-PA

We obtained similar results using CoRe-PA as we did with eMPRess. There exist a negative correlation between cophylogeny precision and average host and symbiont tree topology error. The confidence band around the simple linear regressions were tight, indicating the data points clustered around the regression line.
Contrary to eMPRess results, the mixed-stinkbug model condition obtained nearly horizontal regression line, showing that for this dataset, 15% perturbance in the tree topology did not result in appreciable change to the cophylogenetic precision, which remained low at under 5% cophylogenetic precision. The original annotation cophylogeny reconstruction was estimated using eMPRess, which predicted 5 cospeciations, 5 duplications, and 1 host switch event. On the other hand, CoRe-PA reconstructions on the replicate simulations on average predicted 2 cospeciations and 2 duplications. This may be due to the inherent differences between the algorithms implemented in eMPRess and CoRe-PA.
Two key differences exist between CoRe-PA and eMPRess. First, CoRe-PA generates multiple reconciliations per execution and to limit the number of pairwise comparisons, we limit the maximum number of cophylogeny reconstruction precision calculation to ten per pair of reconstructions. Second, CoRe-PA explores the event penalty space to produce various reconciliations per execution. It is foreseeable for CoRe-PA on small number of taxa datasets like mixed-stinkbug ($n_{host} = 7$, $n_{symbiont} = 12$) to obtain all possible cophylogenetic reconstructions with its various cost schemes per execution. Resulting in a slope of zero for the simple linear regression line. Similar to the mixed-stinkbug model conditions, the mixed-damselfly model condition and the mixed-moth model condition observed their simple linear regression lines' slope to be smaller in magnitude in CoRe-PA results than in eMPRess results.

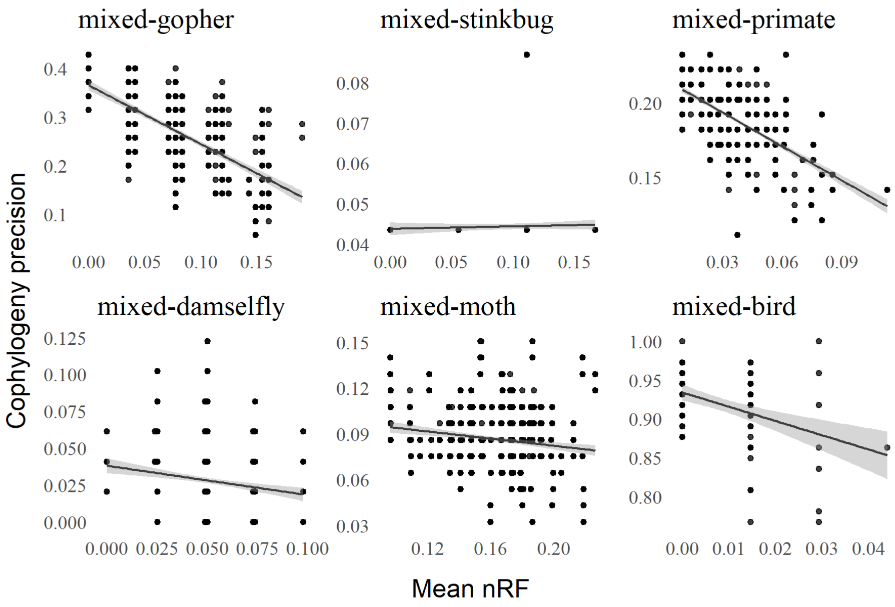

Figure S12: Mixed simulation datasets: precision of CoRe-PA reconciliations compared with averaged host and symbiont tree normalized Robinson-Fould (nRF) distances. For each height scaling factor, a replicate set of 100 alignments were simulated. Co-phylogenetic reconciliation precision was calculated as the aggregate statistic for events found in all of the replicate cophylogeny reconstructions and their respective, original annotation cophylogeny reconstruction.

| | Simple Linear Regression | | | | |
|---|---|---|---|---|---|
| Model conditions | intercept | B coefficient | $R^2$ | RSE | p-value |
| mixed-gopher | 0.3655 | -1.2081 | 0.4621 | 0.0606 | 0.0000 |
| mixed-stinkbug | 0.0438 | 0.0061 | 0.0022 | 0.0061 | 0.0000 |
| mixed-primate | 0.2161 | -0.7561 | 0.3726 | 0.0196 | 0.0000 |
| mixed-damselfly | 0.0381 | -0.1989 | 0.0218 | 0.0264 | 0.0173 |
| mixed-moth | 0.1056 | -0.1167 | 0.0194 | 0.0225 | 0.0000 |
| mixed-bird | 0.9341 | -1.8328 | 0.1663 | 0.0408 | 0.0000 |

Table S3: Simple linear regression details for mixed simulation study evaluated with CoRe-PA.

#### S4.2 Backward-time simulation results with CoRe-PA

In backward-time simulations, we obtained similar results using CoRe-PA as we did with eMPRess such that there exist a negative correlation between cophylogeny precision and average host and symbiont tree topology error. The data points clustered around the regression line as indicated by the tight confidence band around the simple linear regressions line.

| | Simple Linear Regression | | | | |
|---|---|---|---|---|---|
| Model conditions | intercept | B coefficient | $R^2$ | RSE | p-value |
| backward-10 | 0.4689 | -1.6189 | 0.3565 | 0.1031 | 0.0000 |
| backward-50 | 0.4327 | -0.9491 | 0.2813 | 0.0481 | 0.0000 |
| backward-100 | 0.4305 | -2.9033 | 0.3227 | 0.0333 | 0.0000 |
| backward-500 | 0.5380 | -1.7934 | 0.2201 | 0.0210 | 0.0000 |

Table S4: Simple linear regression details for backward-time simulations evaluated with CoRe-PA.

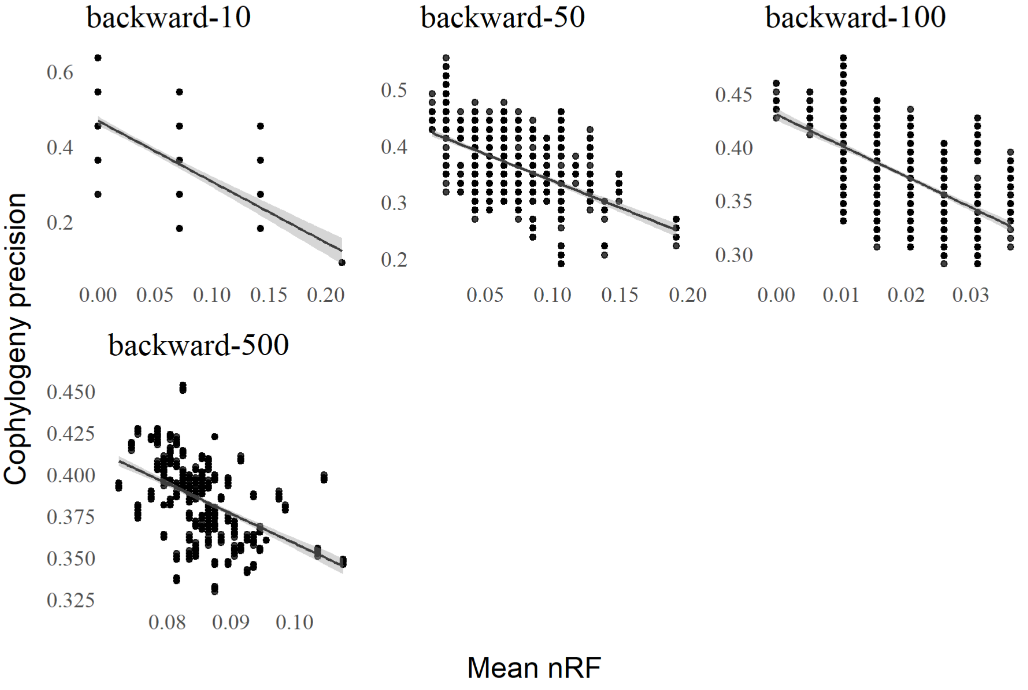

Figure S13: Backward-time simulation datasets: precision of CoRe-PA reconciliations compared with averaged host and symbiont tree normalized Robinson-Fould (nRF) distances. For each height scaling factor, a replicate set of 100 alignments were simulated. Co-phylogenetic reconciliation precision was calculated as the aggregate statistic for events found in all of the replicate cophylogeny reconstructions and their respective, original annotation cophylogeny reconstruction.

##### S4.3 Forward-time simulation results with CoRe-PA

In forward-time simulations, we obtained similar results using CoRe-PA as we did with eMPRess. We found a negative correlation between cophylogeny precision and average host and symbiont tree topology error. The confidence band around the simple linear regressions were tight, indicating the data points clustered around the regression line. The forward-damselfly model condition corresponded with the mixed-damselfly model condition in mixed simulations, which also demonstrated a linear regression line slope that was smaller in magnitude in CoRe-PA results than in eMPRess results. Similarly, forward-bird model condition corresponded with the mixed-bird model condition in mixed simulations, and it also demonstrated a linear regression line slope that was smaller in magnitude in CoRe-PA results than in eMPRess results. Contrary to mixed simulations, forward-stinkbug simulated the mixed-stinkbug model condition but instead of obtaining a horizontal regression slope, forward-stinkbug obtained a trendline closer to model conditions mixed-stinkbug and forward-stinkbug from eMPRess results. This result may support our previous analysis that mixed-stinkbug contained few extant taxa , leading CoRe-PA's multiple reconciliations per execution method to output nearly all possible reconciliations each run. forward-stinkbug model condition ($n_{host} = 16$, $n_{symbiont} = 14$) mimicked mixed-stinkbug model condition ($n_{host} = 7$, $n_{symbiont} = 12$) imperfectly, simulating more than double the number of hosts, which may result in more possible cophylogeny reconstructions since there were more hosts available for the symbionts to interact with and potentially spawn more diverse cophylogenetic event histories.

| | Simple Linear Regression | | | | |
|---|---|---|---|---|---|
| Model conditions | intercept | B coefficient | $R^2$ | RSE | p-value |
| forward-gopher | 0.6913 | -1.0173 | 0.4635 | 0.0707 | 0.0000 |
| forward-stinkbug | 0.6470 | -1.1315 | 0.5401 | 0.0641 | 0.0000 |
| forward-primate | 0.4654 | -0.9690 | 0.7348 | 0.0315 | 0.0000 |
| forward-damselfly | 0.1813 | 0.0374 | 0.0014 | 0.0346 | 0.3090 |
| forward-bird | 0.2309 | -0.4118 | 0.1380 | 0.0230 | 0.0000 |

Table S5: Simple linear regression details for forward-time simulations evaluated with CoRe-PA.

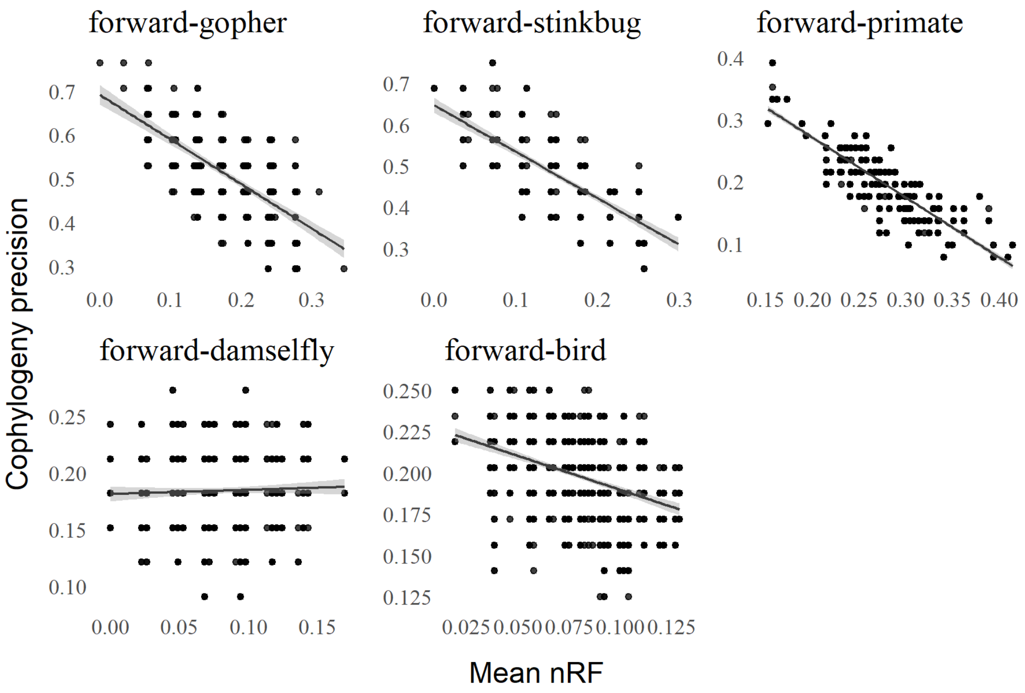

Figure S14: Forward-time simulation datasets: accuracy of CoRe-PA reconciliations compared with averaged host and symbiont tree normalized Robinson-Fould (nRF) distances. Co-phylogenetic reconciliation accuracy was calculated as the aggregate statistic for events found in the 100 replicate cophylogeny reconstructions that were also found in the true coevolutionary history.

### S5 Commands to run cophylogenetic reconciliation software

EMPRess v1.2.1 [Santichaivekin et al., 2021] was used to reconcile cophylogenies in two ways. First, we ran eMPRess v1.2.1 with default cost scheme.

```
python empress_cli.py reconcile {host tree file} {symbiont tree file}
{extant species associations} --csv {out file name}.csv
```

Second, we ran eMPRess v1.2.1 with modified event cost schemes.

```
python empress_cli.py reconcile {host tree file} {symbiont tree file}
{extant species associations} {event cost frequencies} --csv {out file}.csv
```

CoRe-PA version 0.5.2 [Merkle et al., 2010] was used to reconcile cophylogenies and to generate alternative event cost schemes.

```
java -jar core-pa_cli_0.5.2.jar -i {CoRe-PA's nexus format file} -o {out file}
```

COALA version 1.2.1 [Baudet et al., 2015] was used to calculate alternative event cost schemes.

```
java -Xms4056M -Xms8g -jar Coala-1.2.1.jar -input {nexus format file}
-cluster -threads 16
```

### S6 Commands used in empirical experiments

Note that texts inside curly brackets {} indicate files and inputs the user passes into the software, thus they are not part of the command.
BBtools version 37.62 [Bushnell, 2018] was invoked to run BBMap, BBDuk, and Reformat. The following BBDuk command was used to filter and trim Illumina short reads to reduce artifacts and contaminants.

```
# run if lanes 1 and 2 are separate files
bbduk.sh in1={lane1 reads} in2={lane2 reads} out1={paired reads 1}
    out2={paired reads 2} ref=bbmap_adaptor.fa forcetrimleft=5 minlen=90

# run if you have interleaved reads
```

```
bbduk.sh in={interleaved reads} out1={lane1 reads} out2={lane2 reads}
reformat.sh in1={lane1 reads} in2={lane2 reads} out1={paired reads 1}
    out2={paired reads 2} ref=bbmap_adaptor.fa forcetrimleft=5 minlen=90

# produce summary statistics for assembly
statswrapper.sh {assembly} format=4 >> {out file}
```

SPAdes version 3.15.5 [Bankevich et al., 2012] was used to assemble paired short reads.

```
spades.py -k 21,33,55,77,99,127 -o {directory} -1 {paired reads 1}
    -2 {paired reads 2} -t 16
```

BUSCO version 5.3.2 [Simão et al., 2015] was used to assess the completeness of the assemblies.

```
busco -i $fungi -l burkholderiales_odb10 -o {out directory} -m genome -c 4
    --force #bacteria
busco -i $endobac -l mucoromycota_odb10 -o {out directory} -m genome -c 4
    --force #fungi
```

CANU version 2.2 [Koren et al., 2017] was used to assemble PacBio long reads.

```
canu -p {assembly prefix} -d {directory} genomeSize={size in bases} -pacbio {pacbio reads}
```

BLAST+ version 2.2.31 was used to query *Mortierella spp.* and endobacterial assembled contigs from their respective *de novo* assemblies. Seqtk version 1.3 was used to extract contigs from assembly using the blasted bed file to produce fasta format contigs.

```
blastn -query {assembly} -outfmt 6 -max_target_seqs 200 -db {reference} -out {blast file}
awk '!_[$1]++' {blast file} > {bed file}
seqtk subseq -l 60 {blast file} {bed file} > {fasta file}
```

MUMmer version 3.23 [Delcher et al., 2003] was used to variant call the extracted *Mortierella spp.* and endobacterial contigs against their respective reference genomes. SAMtools version 1.15 was used to index and retrieve the VCF file.

```
nucmer --prefix={prefix name} {blasted contigs} {reference genome}
show-snps -Clr -x 1 -T {SNPs prefix}.delta > {SNPs prefix}.snps
MUMmerSNPs2VCF.py {SNPs prefix}.snps {SNPs prefix}.vcf
bgzip -c {SNPs prefix}.vcf > {SNPs prefix}.vcf.gz
tabix -p vcf {SNPs prefix}.vcf.gz
```

Barrnap version 0.9 [Seemann, 2018] was used to extract rRNA genes from *Mortierella spp.* assembly.

```
barrnap --kingdom euk --threads 8 -o {out directory} < {assembly} > {extract rRNA genes}
```

PROKKA version 1.14.6 [Seemann, 2014] was used to extract rRNA genes from *Mortierella's* endobacterial assembly.

```
prokka {assembly} --centre X --compliant --force
```

RAxML version 8.2.12 [Stamatakis, 2014] was used to reconstruct phylogenies under specified software (GTR, HKY85, JC69, and K80).

```
raxmlHPC -m GTRGAMMA -s {unrooted tree} --{software} -p {random number}
-n {out file suffix}
```

RAxML version 8.2.12 [Stamatakis, 2014] was used to bootstrap alignments.

```
raxmlHPC -f j -b {random number} -# {number of samples} -m GTRGAMMA
-s {alignment} -n {out file suffix}
```

RAxML version 8.2.12 [Stamatakis, 2014] was used to midpoint root the phylogenies.

```
raxmlHPC -f I -m GTRCAT -t {unrooted tree} -n {rooted tree file suffix}
-p {random number}
```

PAUP* 4.0 [Swofford, 2003] was used to reconstruct phylogenies under NJ, UPGMA, and SVDquartet.

```
paup4a168_centos64
exe {alignment file}
{lower case model name}
savetree file={out tree file} brlen=yes
quit
```

### S7 Commands used in simulation experiments

Note that texts inside curly brackets {} indicate files and inputs the user passes into the software, thus they are not part of the command.
MAFFT v7.490 [Katoh and Standley, 2013] was used to align sequences in empirical datasets that provided unaligned sequence data.

```
mafft {unaligned sequence file} > {alignment file}
```

Seq-Gen v1.3.4 [Rambaut and Grass, 1997] was used to simulate gap-less alignments under model species trees.

```
seq-gen -mGTR -r{GTR rate parameters} -z {random number} -or
    -l{simulated alignment length} -f{nucleotide frequencies}
    < {model species tree file} > {simulated alignment file}
```

RAxML version 8.2.12 [Stamatakis, 2014] was used to reconstruct phylogenies under the GTR model.

```
raxmlHPC -m GTRGAMMA -s {alignment file} -p {random number} -n {tree file suffix}
```

RAxML version 8.2.12 [Stamatakis, 2014] was used to midpoint root the phylogenies.

```
raxmlHPC -f I -m GTRCAT -t {unrooted tree} -n {rooted tree file suffix}
-p {random number}
```

INDELible version 1.03 [Fletcher and Yang, 2009] was used to simulate n-taxa trees that serve as input to reverse-time simulator originally from [Avino et al., 2019]. To run INDELible, use the following command in the same folder as a INDELible control file called "control.txt".

```
indelible
```

We used the following code in INDELible control file to sample an $n$-taxa tree topology under a birth-death model with birth rate 2.4, death rate 1.1, sampling fraction 0.2566, and mutation rate 0.34.

```
[TYPE] NUCLEOTIDE 1
[TREE] tree1
  [unrooted] 10 2.4 1.1 0.2566 0.34
```

We used the following code in INDELible control file to assign branch lengths using the GTR parameter rates and nucleotide frequencies from the original annotation of the empirical dataset [de Moya et al., 2019] on avian feather lice.

```
[TYPE] NUCLEOTIDE 1
[MODEL] GTRmodel
  [submodel] GTR 1.475477 4.831617 1.410614 1.732842 7.069432
  [statefreq] 0.319 0.192 0.223 0.266
[TREE] tree1 {newick format tree topology from previous INDELible step}
  [branchlengths] NON-ULTRAMETRIC
[PARTITIONS] taxapartition
  [tree1 GTRmodel 1000]
[EVOLVE] taxapartition 1 species_tree
```

### S8 Commands to run simulator software

A modified version of the reverse-time nested coalescent simulator by [Avino et al., 2019] was used to simulate host tree, symbiont tree, and output the true coevolutionary history. To the best of our knowledge, this simulator was not published under copyleft license, therefore we did not include the modified scripts used in this performance study. The following command was used to run the original reverse-time cophylogeny simulator.

```
python nestedCoalescent.py {rooted host tree file} 0.8 0.3 0.4 {symbiont tree file}
```

Treeducken v1.1.0 [Dismukes and Heath, 2021] R software was used to simulate the host tree, the symbiont tree, and the extant species associations. We modified Treeducken data structures to additionally output the true coevolutionary history for the pair of trees in the next section. The following R code was used to run Treeducken v1.1.0 software.

```
library(treeducken)
lambda_H <- {see Treeducken parameters table}
mu_H <- {see Treeducken parameters table}
lambda_C <- {see Treeducken parameters table}
lambda_S <- {see Treeducken parameters table}
mu_S <- {see Treeducken parameters table}
time <- {see Treeducken parameters table}
cophy_obj <- sim_cophylo_bdp(hbr = lambda_H,
                                hdr = mu_H,
                                sbr = lambda_S,
                                sdr = mu_S,
                                cosp_rate =lambda_C,
                                host_exp_rate = 0.0,
                                time_to_sim = time,
                                numbsim = 1)
```

#### S9 Custom-modified Treeducken code

The following R code was used to modify Treeducken's data structures post simulation to rename coevolution events and output the desired format trees with internal node labeling as well as the true, coevolutionary history.

```
library(treeducken)
library(ape)
library(geiger)

# Run Treeducken as normal
lambda_H <- {see Treeducken parameters table}
mu_H <- {see Treeducken parameters table}
lambda_C <- {see Treeducken parameters table}
lambda_S <- {see Treeducken parameters table}
mu_S <- {see Treeducken parameters table}
time <- {see Treeducken parameters table}
cophy_obj <- sim_cophylo_bdp(hbr = lambda_H,
                                hdr = mu_H,
                                sbr = lambda_S,
                                sdr = mu_S,
                                cosp_rate =lambda_C,
                                host_exp_rate = 0.0,
                                time_to_sim = time,
                                numbsim = 1)

# Start modifying phylo and associations data objects
# to output the coevolutionary history with the event types we want

### label internal nodes ###
label_internal_nodes <- function(tree){ #where tree is a phylo object
  tot_internal_nodes<-tree$Nnode # total number of nodes
  start_internal_nodes<-length(tree$tip.label)+1
  end_internal_nodes<-start_internal_nodes+tot_internal_nodes-1
  labels<-list()
  for (i in start_internal_nodes:end_internal_nodes){
    # nodes start incrementing from number of tips
    name<-paste(tips(tree,i),collapse = "_")
    labels <- append(labels, name)
  }
  tree$node.label <- labels
  new_tree <- write.tree(tree)
  return(new_tree)
}
output_unlabeled_tree<-function(tree){
  print(tree)
```

```
  new_tree <- write.tree(tree)
  return(new_tree)
}
#host
write.table(output_unlabeled_tree(cophy_obj[[1]]$host_tree), file_host,
            append = FALSE, sep = " ",
            row.names = FALSE, col.names = FALSE,
            quote=FALSE)
write.table(label_internal_nodes(cophy_obj[[1]]$host_tree), file_host_labeled,
            append = FALSE, sep = " ",
            row.names = FALSE, col.names = FALSE,
            quote=FALSE)
#symb
write.table(output_unlabeled_tree(cophy_obj[[1]]$symb_tree), file_symb,
            append = FALSE, sep = " ",
            row.names = FALSE, col.names = FALSE,
            quote=FALSE)
write.table(label_internal_nodes(cophy_obj[[1]]$symb_tree), file_symb_labeled,
            append = FALSE, sep = " ",
            row.names = FALSE, col.names = FALSE,
            quote=FALSE)

### relabel event history to format: event host_node symb_node) ###
#where tree is a phylo object
relabel_treeducken_event_history <- function(event_history, hosttree, symbtree){
  #host trees
  tot_internal_nodes_h<-hosttree$Nnode # total number of nodes
  num_leaf_host<-length(hosttree$tip.label)
  start_internal_nodes_h<-num_leaf_host+1
  end_internal_nodes_h<-start_internal_nodes_h+tot_internal_nodes_h-1
  labels_host<-list()
  for (i in start_internal_nodes_h:end_internal_nodes_h){
    # nodes start incrementing from number of tips
    name<-paste(tips(hosttree,i),collapse = "_")
    labels_host <- c(labels_host, name)
  }
  hosttree$node.label <- labels_host
  #symb trees
  tot_internal_nodes_s<-symbtree$Nnode # total number of nodes
  num_leaf_symb<-length(symbtree$tip.label)
  start_internal_nodes_s<-num_leaf_symb+1
  end_internal_nodes_s<-start_internal_nodes_s+tot_internal_nodes_s-1
  labels_symb<-list()
  for (i in start_internal_nodes_s:end_internal_nodes_s){
    # nodes start incrementing from number of tips
    name<-paste(tips(symbtree,i),collapse = "_")
    labels_symb <- c(labels_symb, name)
  }
  symbtree$node.label <- labels_symb
  num_events<-nrow(event_history)
  events<-c()
  hosts<-c()
  symbs<-c()
  prefix_host<-"H"  # H for host, S for symb
  prefix_symb<-"S"

  # update event names in Treeducken to the known 4 events that works with cophy software
  # https://github.com/wadedismukes/treeducken/blob/main/src/Simulator.cpp#L682
  treeducken_events=c("SX", "HX", "SSP", "HSP", "AG", "AL", "CSP", "DISP","EXTP", "SHE", "SHS")
  known_events=c("loss", "loss", "duplication", "host_switch",
              "duplication", "loss", "cospeciation", "cospeciation",
```

```
                "loss", "host_switch","host_Switch")
  event_renaming=data.frame(treeducken_events, known_events)
  # mapping to known format event history
  for (i in 1:num_events){
    print(i)
    if (event_history$Event_Type[i] == "I"){
      print("Initialized")
      #skip this one, "I" stands for initialize event vector.
      next
    }
    else{
      new_event<-event_renaming$known_events[event_renaming$treeducken_events
                                          ==event_history$Event_Type[i]]
      events <- c(events, new_event) # events
    }
    if (event_history$Host_Index[i] > num_leaf_host){ #hosts
      hosts <- c(hosts, labels_host[event_history$Host_Index[i]-num_leaf_host])
    }
    else{
      hosts <- c(hosts, paste0(prefix_host,event_history$Host_Index[i]))
    }
    if (event_history$Symbiont_Index[i] > num_leaf_symb){ #symbs
      symbs <- c(symbs, labels_symb[event_history$Symbiont_Index[i]-num_leaf_symb])
    }
    else{
      symbs <- c(symbs, paste0(prefix_symb,event_history$Symbiont_Index[i]))
    }
  }
  new_event_history<-data.frame(events, paste(hosts, sep=" "),
            data.frame("symbs" = paste(symbs, sep=" ")))
            colnames(new_event_history) <- c("events", "hosts", "symbs")
  print(new_event_history)
  return(new_event_history)
}
new_event_history<-relabel_treeducken_event_history(cophy_obj[[1]]$event_history,
            cophy_obj[[1]]$host_tree, cophy_obj[[1]]$symb_tree)
write.table(new_event_history, file_event_history,
            append = FALSE, sep = " ",
            row.names = FALSE, col.names = FALSE,
            quote=FALSE)
### output nexus and empress association links ###
Which.names <- function(DF, value, file_empress_link, file_nexus_link){
  ind <- as.data.frame(which(DF==value, arr.ind=TRUE, useNames =TRUE))
  print(ind)
  num_links<-length(colnames(DF))
  links_empress<-""
  links_nexus<-""
  for (i in 1:num_links){
    symb<-colnames(association_mat)[ind$col[i]]
    host<-rownames(association_mat)[ind$row[i]]
    links_empress<-paste(links_empress,paste(symb, host ,sep=":"), sep="\n")
    links_nexus<-paste(links_nexus,paste0("'",symb,"':'",host,"',",collapse=""), sep="\n")
  }
  links_empress<-sub(".", "", links_empress) # remove first character \n
  links_nexus<-sub(".", "", links_nexus)
  cat(links_empress)
  links_nexus <- gsub(".{1}$", ";",links_nexus) # replace last character with ";"
  cat(links_nexus)
  write(links_empress, file_empress_link)
  write(links_nexus, file_nexus_link)
}
```

```
association_mat<-cophy_obj[[1]]$association_mat
# where cell value is 1 means association exists
Which.names(association_mat, 1, links_empress, links_nexus)
cophy_obj[[1]]$host_tree$Nnode
cophy_obj[[1]]$symb_tree$Nnode
length(new_event_history$events)
sum(new_event_history$events=="cospeciation")
sum(new_event_history$events=="duplication")
sum(new_event_history$events=="host_switch")
num_links<-length(colnames(association_mat))
ind <- as.data.frame(which(association_mat==1, arr.ind=TRUE, useNames =TRUE))
all_symb=c()
# the following only matters if the cophylogenetic software doesn't allow
# a symbiont to associate with multiple hosts. eMPRess and CoRe-PA don't mind.
for (i in 1:num_links){
  symb<-colnames(association_mat)[ind$col[i]]
  if(sum(all_symb==symb) < 1){
    all_symb<-append(all_symb,symb)
  }
  else{
    print("symb lineage on multiple hosts.")
    break
  }
}
```
